# The scaffold nucleoporin Nup188 undergoes liquid-liquid phase separation

**DOI:** 10.64898/2026.10.06.757041

**Authors:** L B Aswathy, Bhawna Burdak, Shrankhla Bawaria, Subhajit Das, Sejal Agarwal, Carl M. Jones, Akshaykumar B. Chaudhari, Vidisha Tripathi, Kishal Roy, Atharv Tiwatane, Parashuram Sonawane, Shekhar C. Mande, Jomon Joseph, Sushma N. Grellscheid, Radha Chauhan

**Affiliations:** Biotechnology Research and Innovation Council - National Centre for Cell Science (BRIC-NCCS), S.P. Pune University Campus, Pune, Maharashtra, 411007, India; Regional Centre for Biotechnology (RCB), Faridabad; Computational Biology Unit, Department of Informatics, University of Bergen, Bergen, 5006, Norway; Department of Bioinformatics, Savitribai Phule Pune University, Pune, India 411007; ImmunitoAI Pvt. Ltd., Bengaluru, India; Architecture et Fonction des Macromolécules Biologiques (AFMB), UMR7257 Aix-Marseille University, 13288 Marseille, CEDEX 09 France; Department of Biological Sciences, Indian Institute of Science Education and Research (IISER), Mohali, Punjab, India; Department of Biomedicine, University of Bergen, Bergen, 5009, Norway; Department of Biosciences, University of Durham, Durham, DH1 3LE, United Kingdom of Great Britain and Northern Ireland (the); Vollum institute, Oregon health and science university, Portland Oregon USA 97239

**Keywords:** Liquid-Liquid Phase Separation, hNup188, IDR

## Abstract

Components of nuclear pore complexes are broadly classified into two categories: intrinsically disordered FG-repeat nucleoporins such as Nup98, which can undergo liquid–liquid phase separation, and scaffold nucleoporins, which form the NPC’s structural framework. The basis for crosstalk of these nucleoporins within NPC and beyond is poorly understood. Here we show that human Nup188, a helical inner-ring scaffold protein, forms condensates both in purified form as well as at endogenous level. This condensate exhibits fusion, dissolved in 1,6-hexanediol and recovered after photobleaching, with dynamics and material properties comparable to those of hNup98 condensates. Moreover, these condensates are not colocalized to annulate lamellae. Our analysis also revealed that hNup188CTD is sufficient to form condensates and MD simulations revealed the key roles of IDR1–3 for condensation. Notably, hNup93 expression along with hNup188 suppressed the condensate formation. Together, these findings indicate that a structured scaffold nucleoporin can adopt a condensate-forming state via its CTD and Nup93 binding can suppress this phenomenon.

## Introduction

Nuclear pore complexes (NPCs) are multi-protein assemblies of roughly 120 MDa that span the nuclear envelope and regulate transport between the nucleus and the cytoplasm (Beck & Hurt, 2017; Kosinski et al., 2016; Obarska-Kosinska et al., 2026). Each NPC is built from multiple copies of approximately 30 different nucleoporins (Nups) that assemble into subcomplexes around an eightfold-symmetric core (Beck & Hurt, 2017; Knockenhauer & Schwartz, 2016; Mosalaganti et al., 2022). Among the subcomplexes, the central transport channel (CTC) has been studied in detail because of its direct role in nuclear transport (Chug et al., 2015; Schrader et al., 2008; Sharma et al., 2015; Sonawane et al., 2020; Ulrich et al., 2014). In contrast, the inner ring complex (IRC), also called the scaffold nucleoporin complex, has been explored far less, largely due to its challenging size and difficulty in purification. Integrated structural studies show that the IRC links the CTC to the Nups embedded in the nuclear envelope and may therefore contribute to the contraction and dilation of the NPC (Akey et al., 2022; Mosalaganti et al., 2022; Zimmerli et al., 2021). In vertebrates, the inner ring is formed mainly by Nup93 complexes, in which the linker Nup93 associates with Nup155, Nup35 (Nup53) and either Nup188 or Nup205 (Vollmer and Antonin, 2014). Studies in Xenopus and yeast showed that Nup188 and its paralog Nup205 bind Nup93 in a mutually exclusive manner (Amlacher et al., 2011; Theerthagiri et al., 2010) and an integrative structural model of the human NPC placed 16 copies of each paralog adjacent to Nup93 in a low-resolution NPC density map (Mosalaganti et al., 2022). High resolution structures of fungal homologs of Nup188 and Nup205 showed that these are α-solenoid proteins whose S-like architecture resembles nuclear transport receptors (NTRs or karyopherins) (Andersen et al., 2013; Flemming et al., 2012; Sampathkumar et al., 2013; Stuwe et al., 2014). Consistent with this similarity, both proteins bind FG repeats, yet the two paralogs are not functionally equivalent (Andersen et al., 2013, Theerthagiri et al., 2010).

In contrast to scaffold Nups, a subset of Nups which lines the centre of NPC, contains intrinsically disordered region rich in Phenylalanine-Glycine repeats called FXFG/GLFG Nucleoporins (Denning et al., 2003; Yu et al., 2023; Devos et al., 2006). These FG-Nups interact multivalently to form a mesh-like network, which creates the dense, phase-separated permeability barrier that is essential for cargo transport across the NPC (Frey et al., 2006; Celetti et al., 2020; Lemke, 2016; Najbauer et al., 2022; Ng & Görlich, 2022; Schmidt & Görlich, 2015) The best-known example is Nup98, whose FG domains are capable of phase separating spontaneously at critical concentrations, and contribute to the selective inclusion properties of NPC (Dekker et al., 2023; Ng & Görlich, 2022; Schmidt & Görlich, 2015). NMR and simulation studies identified that these dynamic FG motifs as the drivers of this cohesion, which is disrupted by 1,6-hexanediol (Dekker et al., 2023; Najbauer et al., 2022; Schmidt & Görlich, 2015).

Liquid-liquid phase separation (LLPS) has emerged as a general mechanism by which cells organize processes such as transcription and DNA repair (Banani et al., 2017; Hnisz et al., 2017; Kilic et al., 2019). Within the NPC, LLPS has been studied mainly for the FG-Nups of the CTC, and cytoplasmic filaments (Celetti et al., 2020; Frey et al., 2006; Ng et al., 2023; Oka et al., 2023; Schmidt & Görlich, 2015). Nucleoporin condensation also contributes to NPC biogenesis: In Drosophila oocytes, nucleoporins condense into granules in which Nup358 plays a key role, and these granules interact and mature into annulate lamellae (AL)-NPCs (Hampoelz et al., 2016, 2019). Although LLPS is usually attributed by Intrinsically Disordered Regions (IDRs), structured and helical elements can also contribute, as in the TDP-43 and in the centrosome scaffold SPD-5, (Conicella et al., 2016; Woodruff et al., 2017). Whether a similarly structured, helical scaffold component of the NPC itself might possess such capacity has not been examined.

Scaffold Nups such as the IRC components are generally shown to provide the structural framework of the NPC. Within the NPC, Nup188 regulates inner nuclear membrane protein transport (Theerthagiri et al., 2010). However, several other observations suggest, Nup188 has additional functions. At the base of cilia, Nup188 and Nup93 are part of two barrel-like structures that encase the centrioles, and their depletion causes cilia loss and abnormal heart looping in Xenopus embryos (del Viso et al., 2016). Besides that, Nup188 is also part of the pericentriolar material and directly binds to Cep152, which is required for centriole duplication (Vishnoi et al., 2020). Together with Nup93 and Nup205, it also contributes to the repression of HOXA genes (Labade et al., 2016). Consistent with these roles, Nup188 is also implicated in human disease. A duplication of Nup188 was found in a patient with heterotaxy and congenital heart disease (Fakhro et al., 2011) whereas biallelic loss-of-function variants of NUP188 cause Sandestig–Stefanova syndrome (Muir et al., 2020; Sandestig et al., 2020; Poot, 2022; Almalki et al., 2026 and Soğukpınar et al., 2026). Heterozygous truncating variants have been linked to mitral valve prolapse (Haskell et al., 2017). Increased Nup188 expression is also reported in different cancers (Zhu et al., 2025). Although these reports highlight the broader clinical and physiological significance of Nup188, its core biochemical properties and their contribution to overall cellular functions remain poorly understood.

Here, we combined computational prediction, *in vitro* assays, cell imaging and molecular dynamics (MD) simulations to establish that human Nup188 (hNup188), an α-helical scaffold protein, can phase separate. We show that hNup188 forms condensates *in vitro* and *in vivo*. Analysis of morphological descriptors and material properties indicates that these condensates resemble those of the FG-Nup Nup98. We identified that the C-terminal region of Nup188 is sufficient for condensation, and simulations show that this region is flexible and phase separate. Finally, we show that binding of Nup93 to Nup188 suppresses condensate formation. Together, our results show that, although hNup188 is a structured helical protein, it can form liquid-like condensates comparable to those of Nup98, and the binding of Nup93 antagonizes this behavior.

## Results

### 1. Human Nup188 sequence harbors features of liquid–liquid phase separation

The human Nup188 (hNup188) was expressed as a C-terminal GFP fusion (Figure 1A) in HEK293F cells and purified using the combination of GFP affinity and size exclusion chromatography (SEC) as described in Supplementary Figure S1. The SEC profile revealed that hNup188-GFP protein eluted as a single peak at 13.8 mL from a Superose6 increase 10/300 column (Figure 1B) and migrated as a single band near to 250 kDa on SDS–PAGE (Figure 1C), consistent with its theoretical molecular weight (226 kDa). Repeated attempts to concentrate the purified protein led to visible precipitation (data not shown), suggesting poor intrinsic solubility or concentration dependent self-association. This observation leads us to examine the phase separation propensity of hNup188.

**Figure 1.**
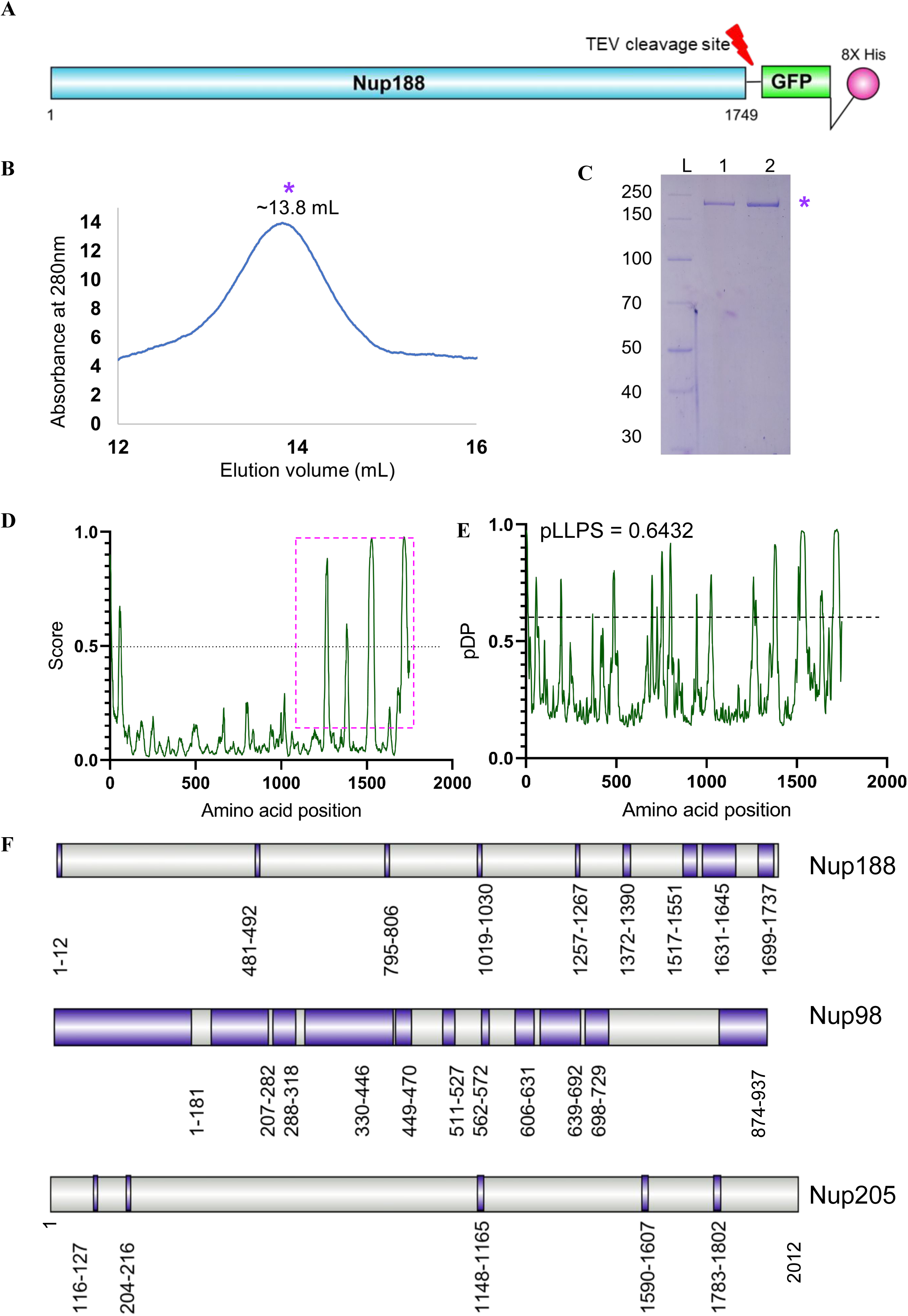
Biophysical and computational characterization of human Nup188. (A) Construct used for expression: human Nup188 (residues 1–1749) followed by a TEV protease cleavage site, GFP, and an 8×His tag at the C-terminus. (B) Size-exclusion chromatography of purified Nup188 on a Superose 6 Increase 10/300 GL column in DDM-containing buffer. Absorbance at 280 nm is plotted against elution volume; the main peak (purple asterisk, ∼13.8 mL) is consistent with a monomeric form. (C) Coomassie-stained SDS-PAGE of the peak fraction (lanes 1 and 2) showing a single band near to molecular weight of ∼250 kDa (*) L, molecular weight marker (kDa). (D) Intrinsic disorder prediction for Nup188 by IUPred3 (Erdős et al., 2021). The dotted line marks the disorder threshold (score = 0.5); the dashed box highlights multiple disordered segments in the C-terminal region. (E) Droplet-promoting propensity (pDP) along the Nup188 sequence predicted by FuzDrop (Hatos et al., 2022). Regions of ≥10 consecutive residues above the threshold (dashed line, 0.6) are predicted to promote droplet formation. The overall liquid–liquid phase separation probability (pLLPS) is 0.6432, indicating a droplet-forming protein. (F) Droplet-promoting regions (purple) of Nup188 (1–1749), Nup98 (1–937; positive control) and Nup205 (1–2012; negative control), as predicted by FuzDrop. Residue ranges are indicated below each region.

Initially, we performed IUPred3 analysis (Erdos et al., 2021), which revealed the presence of intrinsically disordered stretches in the C-terminal region (Figure 1D). Because intrinsically disordered regions (IDRs) frequently promote condensate formation, we next subjected it to the machine-learning predictor FuzDrop (Hatos et al., 2022). In FuzDrop, hNup188 obtained a droplet-formation probability score of 0.6432, which was above the threshold (score = 0.6), and its droplet-promoting stretches predicted to be clustered toward the C-terminus (Figure 1E). For comparison, two other proteins, human Nup98 (hNup98), a predominantly disordered FG-nucleoporin was also analyzed in similar fashion, and served as a positive control (score 1.0), as well as human Nup205 (hNup205), the paralog of Nup188, as a negative control (score 0.2) (Figure 1F). We also applied MolPhase analysis (Liang et al., 2024) which gave concordant results: it predicted disordered stretches at the C-terminus with a score of 0.989 for hNup188 (Supplementary Figure S2A). According to MolPhase, the phase-separation propensity of hNup188 derives mainly from disorder-promoting (57.3%), polar (34.1%), and hydrophobic (38.8%) residues, a profile consistent with established determinants of condensate formation (Supplementary Figure S2B). Together, all these suggested human Nup188 has features essential for condensate formation.

### 2. Purified Nup188 forms condensates *in vitro*

To test experimentally whether hNup188 undergoes phase separation, we performed *in vitro* condensation assays with GFP-tagged hNup188 at 0.2 mg/mL (∼1 μM) at 4 °C in the presence of 1, 2.5 or 5% PEG 5000 as a crowding agent, which is widely utilized as a macromolecular crowding agent to mimic the dense cellular cytoplasm in *in vitro* biophysical and biochemical assays (André et al., 2023; André & Spruijt, 2020) followed by imaging using fluorescence microscopy (488 nm laser). We also imaged the sample after 24 h at 4 °C without a crowding agent. To negate the possible effects of the GFP tag in condensate formation, additionally GFP alone was also assayed in an identical buffer.

Without a crowding agent, hNup188 was homogeneously distributed at the start of the experiment (Supplementary Figure S3A), and a few micron-scale speckles appeared after 24 h at 4 °C (Figure 2A). With 1% PEG 5000, a few droplets appeared after 60 min and remained unchanged at 2 h (Figure 2B). At 2.5% PEG 5000, more droplets formed within 60 min than at 1% (Supplementary Figure S3D), and at 5% PEG 5000 both circular and irregular droplets were visible within 10 min of mixing (Figure 2C). Notably, neither the buffer alone nor the buffer with 10% PEG 5000 showed fluorescence at 488 nm (Supplementary Figure S3B, S3C), excluding buffer autofluorescence. Therefore, it was concluded that across the different PEG concentrations, the droplet formation appeared faster and in greater numbers (Figure 2; Supplementary Figure S3D). Moreover, GFP alone at 1 mg/mL (37 μM), a ∼37-fold higher molar concentration than Nup188, remained homogeneously distributed after 1 h with 2.5, 5 or 10% PEG 5000 (Supplementary Figure S4A, S4B), showing that neither the GFP tag nor the buffer drives droplet formation. Together, these data show that recombinantly purified Nup188 forms condensates in a crowding agent concentration dependent manner. This observation is also in agreement with the computational predictions as described above.

**Figure 2.**
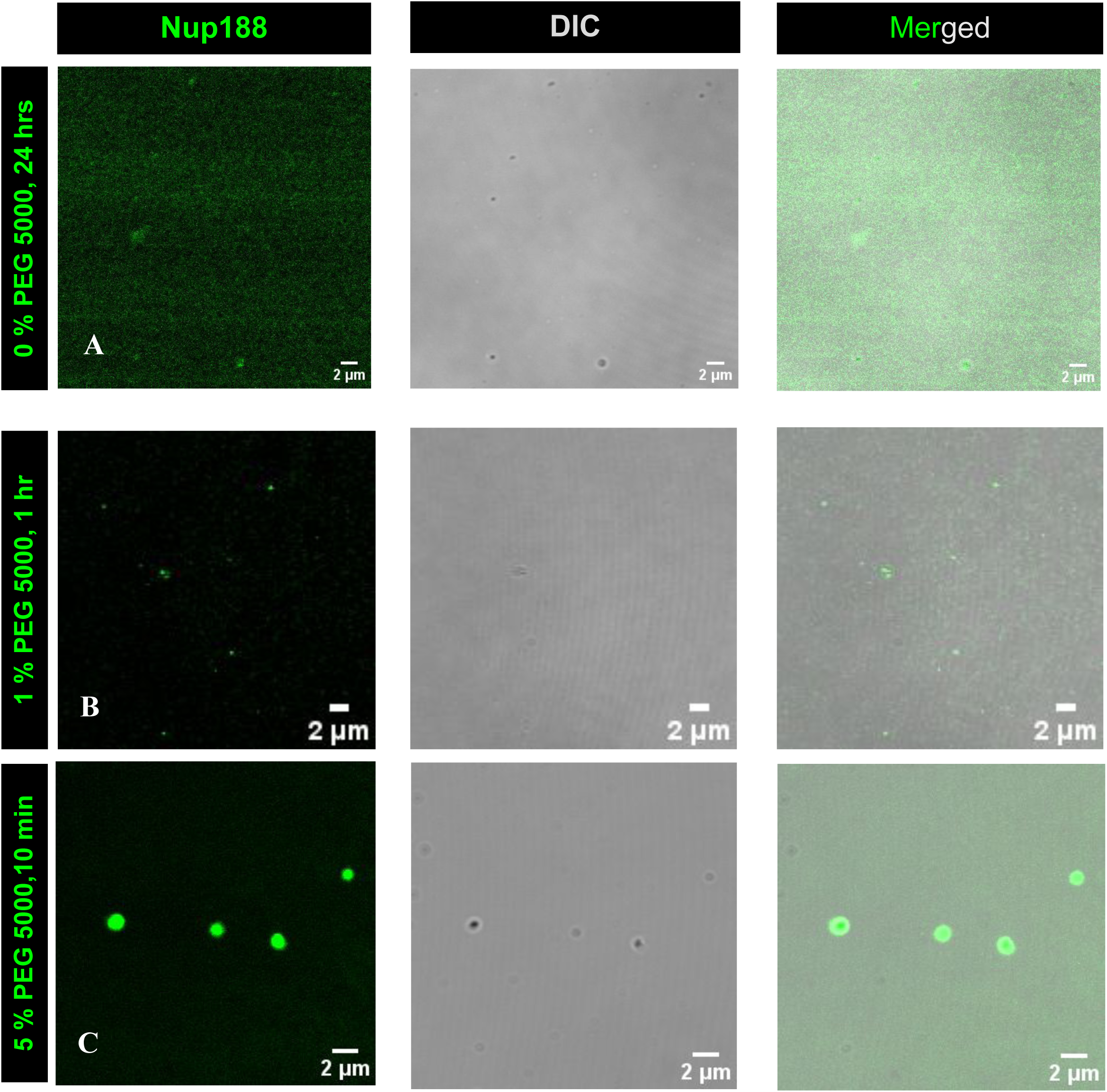
Recombinant Nup188 forms condensates *in vitro*. Fluorescence (Nup188-GFP), DIC and merged images of purified recombinant Nup188 (A) after 24 h at 4 °C without PEG 5000, (B) with 1% PEG 5000 after 1 h, and (C) with 5% PEG 5000 after 10 min. Scale bars, 2 µm. Experiments were repeated independently three times; representative images are shown.

### 3. Endogenous hNup188 localizes to puncta in several cell lines

To assess whether the *in vitro* condensate formation phenomenon of hNup188 has any physiological relevance, we examined the subcellular localization of endogenous hNup188 by immunofluorescence in three different cell lines: WI-38 human lung fibroblasts; the osteosarcoma cancer cell line - U2OS and cervical carcinoma cell line - HeLa. The immunofluorescence analysis of WI-38 fibroblasts (Figure 3A) revealed that hNup188 localized to both cytoplasm and nucleus as small puncta, along with its expected localization at the nuclear envelope (Supplementary Figure S5A). In case of HeLa cells, hNup188 was observed to form a predominant population of puncta inside the nucleus (Figure 3B), however, in case of U2OS cells, these puncta were mostly cytoplasmic, with a few in the nucleus too (Figure 3C). In contrast to the clear nuclear membrane staining in the WI-38 cell line, hNup188 was not observed to be localized in the nuclear envelope of U2OS cells, when co-stained with nuclear envelope marker, mab414 (Supplementary Figure S5B). Altogether, these results suggest that Nup188 indeed forms punctas across various cell-types, but with marked differences in puncta patterns (size and number).

**Figure 3.**
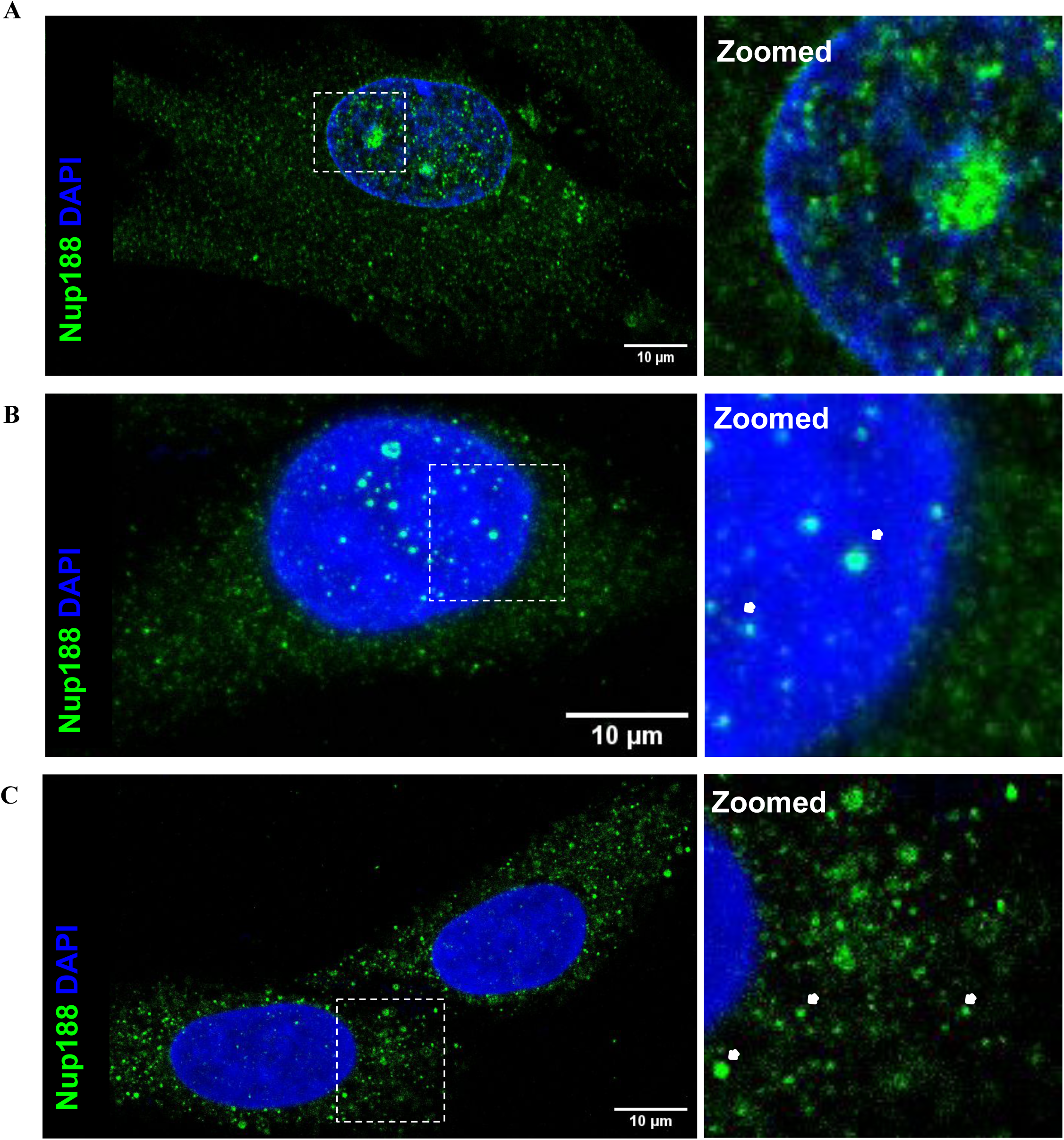
Endogenous Nup188 exhibits puncta distribution in the cell lines. Immunofluorescence of endogenous Nup188 (anti-Nup188, green) with DAPI (blue) in (A) WI-38 human lung fibroblasts, (B) U2OS osteosarcoma cells and (C) HeLa cervical carcinoma cells. Dashed boxes mark the regions enlarged in the right-hand panels; white arrowheads indicate nuclear or cytoplasmic foci. Scale bars, 10 µm.

### 4. Overexpressed hNup188 forms dynamic, liquid-like condensates in cells

To further characterize hNup188 puncta formation, we used an overexpression system (pEG BacMam Nup188-GFP transfected into U2OS/HEK293T cells) to obtain better signals in immunofluorescence assays. Briefly, we transiently transfected the plasmid into U2OS cells and counterstained with DAPI for the genetic material and mab414 for the nuclear membrane (Figure 4A). As expected, the overexpressed Nup188 formed discrete droplets of various sizes inside the cell. This has been observed in other cell types such as HEK293FT also (Supplementary Figure S6A). These observations along with *in vitro* and computational evidence suggested that Nup188 shows LLPS behavior. Finally, more detailed phase-separation analysis was performed by systematically observing droplet fusion and fluorescence recovery after photobleaching (FRAP). These data revealed the individual fusion events, two droplets merged into one that relaxed into a round shape (Movie 2), Such behavior is widely reported for low-viscosity LLPS (Ghosh et al., 2021). Another hallmark feature of such LLPS characteristic is sensitivity to 1, 6 - hexanediol (Ulianov et al., 2021) as LLPS phenomenon occurs due to multivalent weak interactions, 1, 6 - hexanediol can disrupt such interactions. In our study, we observed that Nup188-GFP protein in transfected U2OS cells showed bright discrete droplets in both nucleus and cytoplasm (Figure 4B); however, when such cells were treated with 1, 6 - hexanediol, these droplets dissolved and the fluorescence became diffusely distributed (Figure 4C). Thus, we concluded that hNup188 protein puncta are formed due to liquid-liquid phase separation.

**Figure 4.**
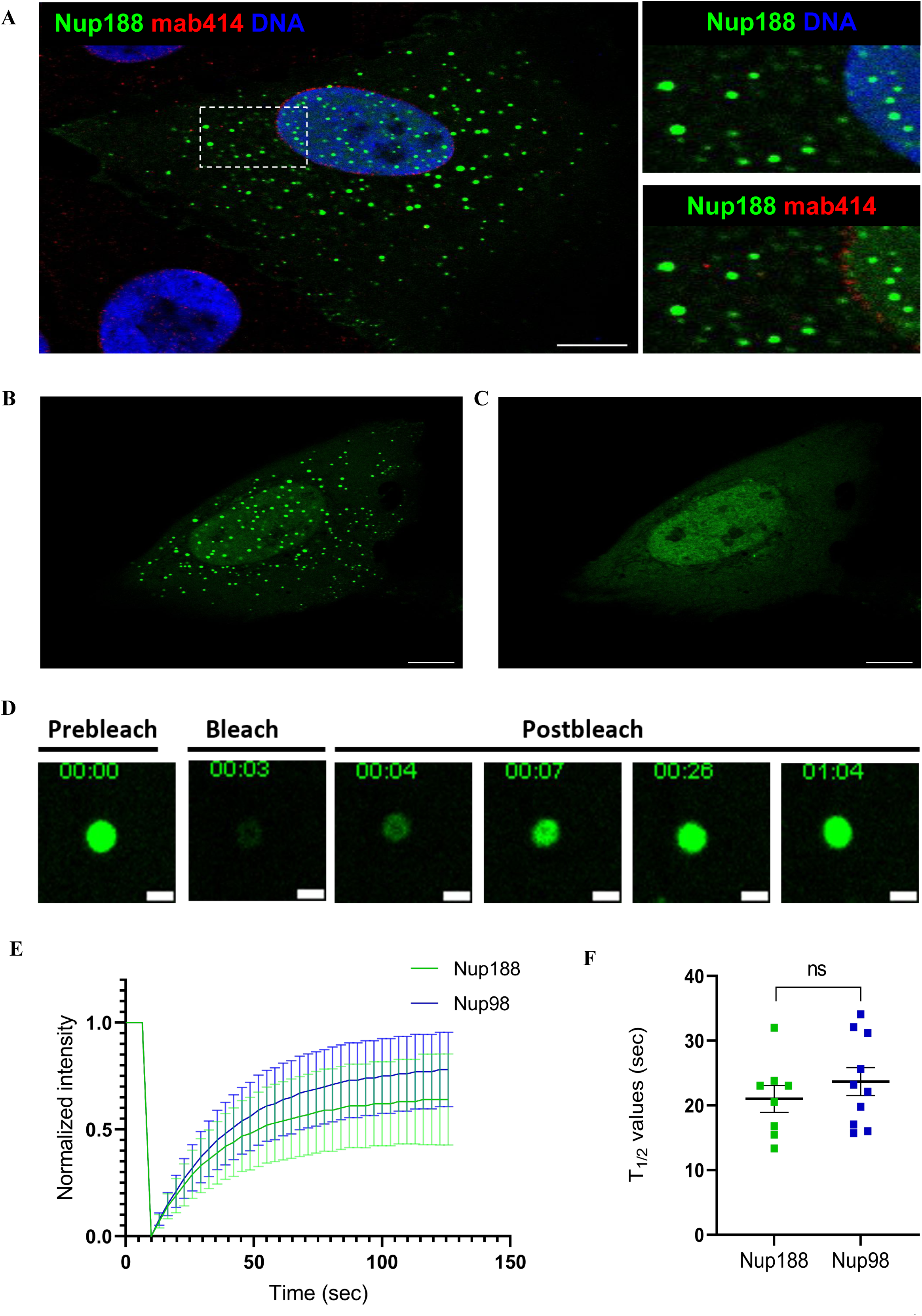
Overexpressed Nup188 forms liquid-like condensates in U2OS cells. (A) Confocal images of U2OS cells transiently expressing Nup188-GFP (green), stained with mab414 (nuclear pore complexes, red) and DAPI (blue). Dashed boxes mark the enlarged regions on the right. Scale bar, 10 µm. Imaging was repeated in three independent experiments (N = 3, n = 10 cells each). (B, C) Nup188-GFP condensates before (B) and 2 s after (C) treatment with 5 % 1,6-hexanediol (n = 7 cells). Scale bars, 10 µm. (D) Representative FRAP of a single Nup188-GFP condensate; time is shown as mm:ss. Scale bar 2 µm (E) Normalized fluorescence recovery after photobleaching (mean ± SD; Nup188, n = 8 condensates; Nup98, n = 10 condensates. FRAP was performed one condensate per cell). Nup98, a known phase-separating FG-nucleoporin, serves as a reference. (F) FRAP half-time (t½) of individual condensates; horizontal bar = median. ns, not significant (two-tailed Mann–Whitney U test).

There are many nucleoporins reported to form LLPS and most of them shown to form condensate due to presence of intrinsically disordered domain as they are enriched with FxFG/FG/GLFG repeats. Among them Nup98 is a widely accepted condensate forming Nucleoporin (Najbauer et al., 2022b; Ng et al., 2023; Schmidt & Görlich, 2015). So, we included the hNup98 condensate formation phenomenon as a reference to further characterize the hNup188 condensates. We performed FRAP analysis on both hNup188-GFP and hNup98-GFP overexpressed protein in U2OS cells. Briefly, the individual droplets/condensates were selected and bleached with 100 % laser intensity and monitored for more than 100s to track the recovery of fluorescence signal. Immediately after bleaching (Figure 4D, Movie 3), both protein condensates showed recovery of fluorescence intensity indicating the LLPS behavior, consistent with the liquid-like mobility inside the condensates (Figure 4E). Moreover, half-time of recovery (t½) did not differ significantly between Nup188 (median = 21.80 s, n = 8) and Nup98 condensates (median = 22.66 s, n = 10) (Mann-Whitney test, exact two-tailed P = 0.4082; Figure 4F). When we compared the mobile fractions, Nup98 (∼0.80) hits slightly higher than Nup188 (∼0.65), which is insignificant (p = 0.0962). This comparable recovery showed that Nup188 condensates possess similar liquid-like properties as GLFG containing Nup98 condensates.

### 5. Nup188 condensates resemble Nup98 condensates in their material properties

Since the FRAP analysis has established that Nup188 exhibits liquid-like behavior, we went on checking whether these condensates also exhibit the mechanical and morphological parameters of classical condensates. We employed Flickerprint, a python package based on flicker spectroscopy that can determine the bending rigidity (κ) and interfacial tension (σ) by analyzing the shape fluctuations of soft bodies in fast recorded live videos on a large scale and non-invasive manner (Law et al., 2023; Williamson et al., 2025). In addition, it also helps to understand the shape descriptors such as circularity, solidity, as well as axis ratios. We recorded fast paced live videos for both hNup98 and hNup188GFP fusion protein condensates in U2OS cells and used them as input for the FlickerPrint analysis as described by Williamson et al., 2025. We recorded a total of 29 and 22 videos for hNup188 and hNup98 respectively. After the final filtering we ended with 7712 (hNup188) and 3018 (hNup98) condensates for the analysis.

Condensate size distribution analysis has shown that the histograms were right skewed, for both Nup188 and Nup98 with the majority population having radii below 0.4 μm and a smaller population extending towards 1.4 μm (Figure 5A & 5B). The graph showed a typical pattern of condensates, where the predominant population is smaller ones and an independent minor subset highlighting bigger ones, which could be the aged ones. In order to understand the mechanical properties of these condensates, we also analyzed condensate radius against interfacial tension (σ) (Figure 5C, D) and bending rigidity (κ) (Figure 5E, 5F). The density scatter plot analysis revealed that the majority of both Nup188 and Nup98 condensates populated a low range of σ (∼10⁻⁷–10⁻⁵ N/m), whereas in case of bending rigidity (κ) values are in broader range (∼10^-2^–10^2^ k_B_T). Both protein condensates showed a major population with low to medium κ values (∼10^-2^–10^1^ k_B_T) whereas a small fraction is also observed with the high κ values (∼10^1^–10^2^ k_B_T). This was consistent with the histogram representation of bending rigidity shown in Supplementary figures (S7A and S7B). Taken together, we observed that both hNup98 and hNup188 condensates have lower interfacial tension/lower bending rigidity which is consistent with liquid-like material behavior, in which condensate boundaries fluctuate readily, and interfacial tension is dominated by surface energy rather than a rigid, elastic boundary. A smaller subset of condensates, generally of larger radius, exhibited higher σ and κ values, suggesting a minority population with reduced fluidity too. The density scatter plot showed a joint relationship between interfacial tension and bending rigidity (Supplementary figure S7C and S7D) further supporting the conclusion that both Nup188, like Nup98, forms predominantly liquid-like condensates.

**Figure 5.**
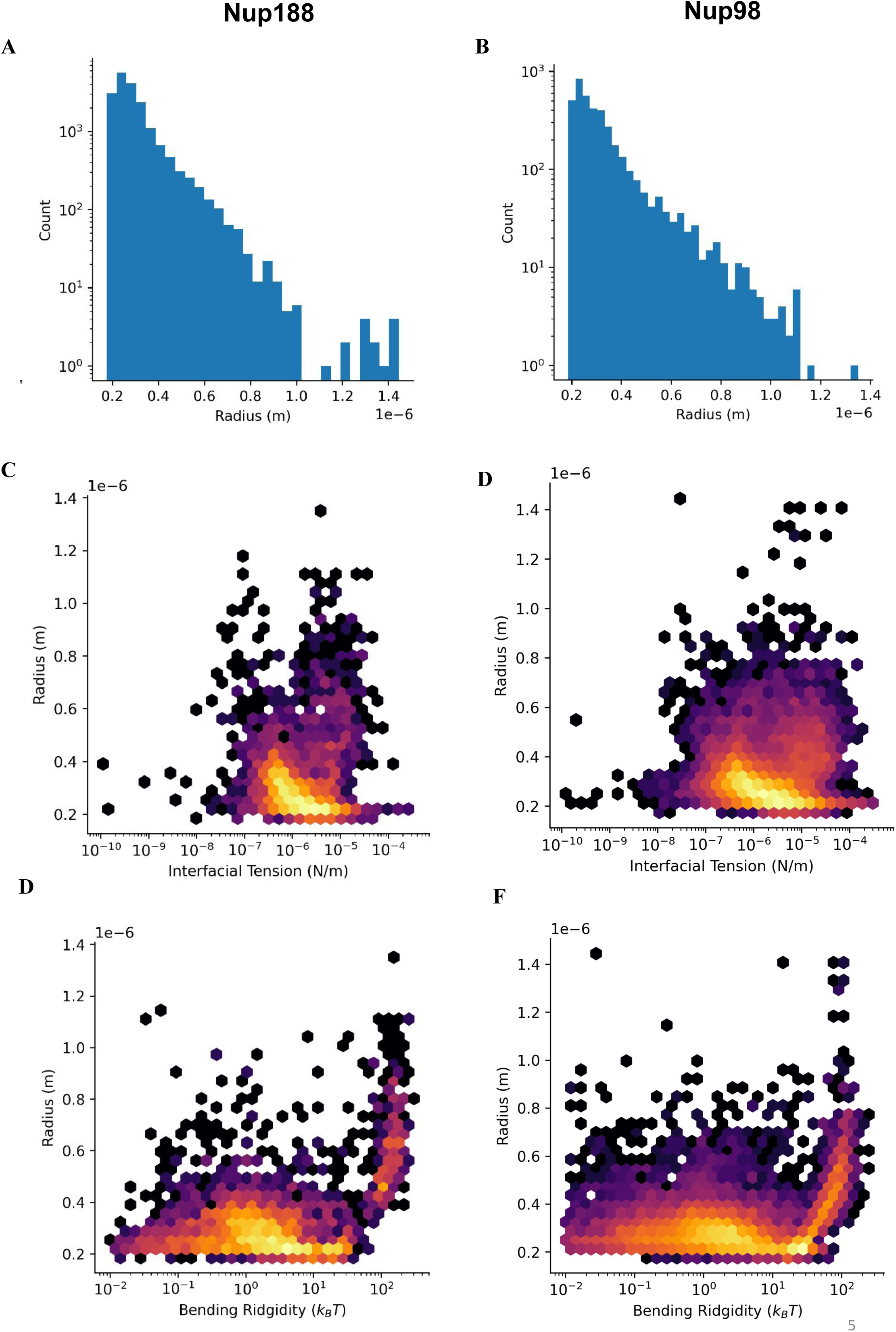
FlickerPrint analysis indicates that material property of Nup188 condensates is similar as Nup98. (A, B) Size distributions (radius) of Nup188 (A) and Nup98 (B) condensates. (C, D) Density scatter plots of condensate radius versus interfacial tension (σ) for Nup188 (C) and Nup98 (D). (E, F) Density scatter plots of radius versus bending rigidity (κ, in k_B_T) for Nup188 (E) and Nup98 (F). Colour indicates local point density (yellow, high; purple, low).

We also analyzed shape descriptors such as circularity, solidity and axis ratio using FlickerPrint (Williamson et al., 2025) to quantitatively verify the fluid phase state of the assemblies (Putnam & Seydoux, 2021; Patel et al., 2015) has reported that isotropic surface tension drives liquid droplets to minimize surface area by adopting spherical geometries. In our study, both hNup98 and hNup188 condensates exhibited circularity skewed toward 1.0, with most condensates between 0.9 and 1.0 (Supplementary Figure S8A, SB), indicating near-circular shapes. Consistent with this, the solidity values, which measure the smooth boundaries - were similarly skewed towards 1.0, further supporting that both Nup188 and Nup98 adopt droplet morphology rather than irregular shapes (Supplementary Figure S8C and S8D). Similarly, axis ratio distributions have shown that the majority of condensates of both proteins had values close to 1.0, showing nearly circular or non-elongated shapes. Altogether, these shape descriptors agree that both Nup188 and Nup98 form condensates with liquid-droplet-like morphology.

### 6. Cytoplasmic Nup188 condensates are distinct from annulate lamellae

Many nucleoporins (Nup358, Nup62 etc.) are known to be localized to the cytoplasm, as they are shown to be resident of annulate lamellae (AL) (Lin et al., 2026; Kessel, 1992; Oliveira and Joseph, 2026). As hNup188 condensates are also present in the cytoplasm (Figure 3), we hypothesized that hNup188 condensate may colocalize to the AL. Using structured illumination microscopy (SIM), we immunostained for the AL marker Nup358 in hNup188–GFP expressing U2OS cells. Surprisingly, we observed that the discrete hNup188 droplets did not colocalize with Nup358 (Figure 6A; Movie 4). We also imaged hNup205, a Nup188 paralog, in a similar experiment and observed that hNup205 colocalized with Nup358 when expressed in U2OS cells (Figure 6B), as reported previously also (Lin et al., 2026). These data indicate that cytoplasmic Nup188 condensates are a distinct pool, separated from ALs, and may therefore have relevance beyond AL-related functions.

**Figure 6.**
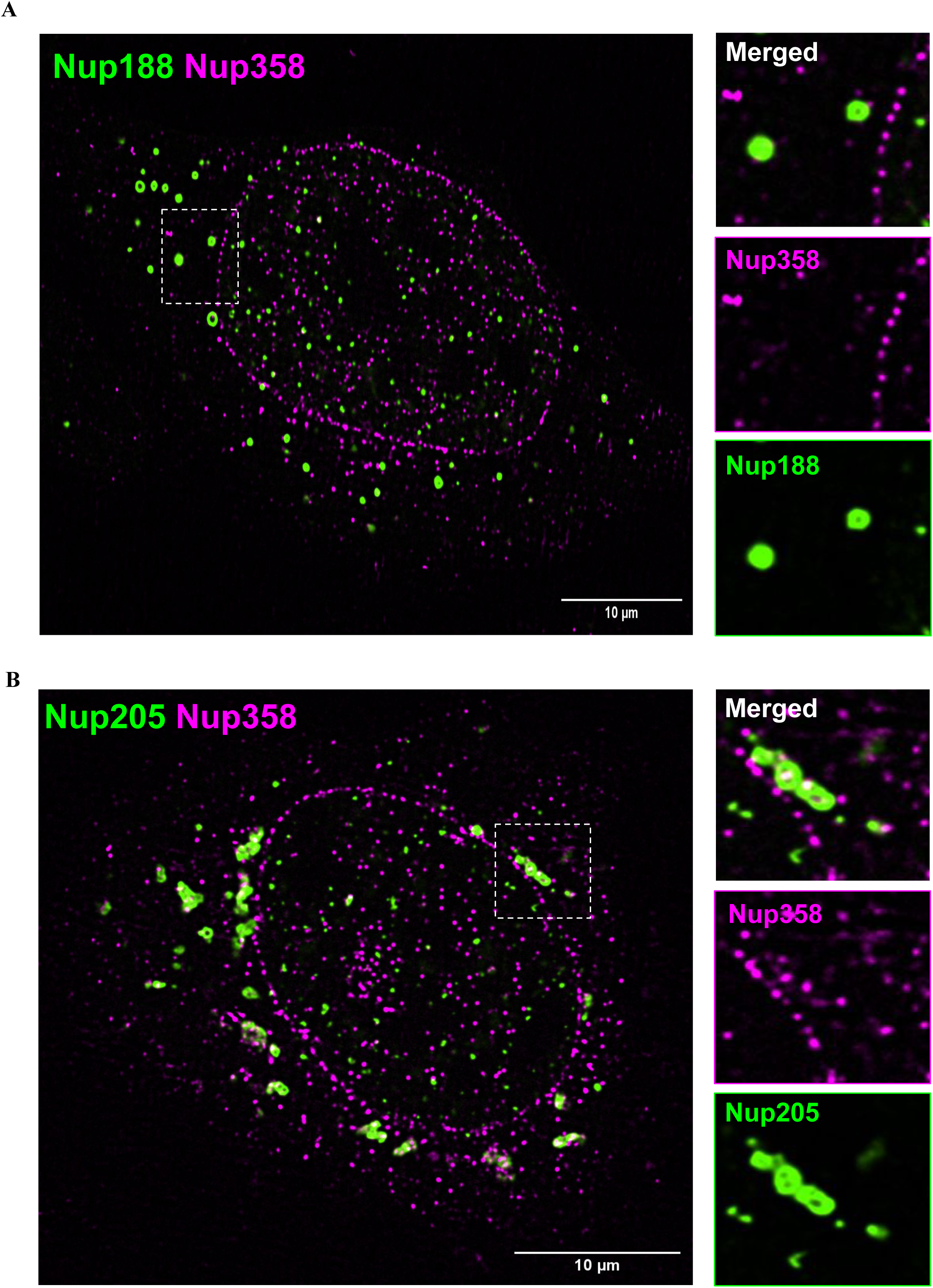
Nup188 condensates are distinct from annulate lamellae. Structured illumination microscopy (SIM) of U2OS cells co-stained for the annulate lamellae marker Nup358 (magenta). (A) Cell expressing Nup188-GFP (green). (B) Cell expressing Nup205-GFP (green). Colocalization appears white. Dashed boxes mark the enlarged regions on the right, which show the merged and individual channels. Scale bars, 10 µm. Imaging was repeated in three independent experiments (N = 3, n = 10 cells each).

### 7. The C-terminal domain of Nup188 is sufficient to form condensates

Our IUPred3 and MolPhase analysis revealed that the C-terminal domain of hNup188 is disordered and has a propensity to form condensates. Therefore, based on PSIPRED (David, 1999) analysis, we divided the whole hNup188 sequence into two parts: N-terminal domain (residues 1–1112; Nup188NTD) and C-terminal domain (residues 1113–1749; Nup188CTD), and prepared expression constructs using the pEGFP-C1 vector backbone (Supplementary Figure S9A) and expressed them in U2OS cells for immunofluorescence analysis. It was observed that the hNup188NTD-GFP fusion protein is diffusely distributed throughout the cell (Figure 7A, i) and no puncta were observed. In contrast, the hNup188CTD-GFP fusion was expressed predominantly in the cytoplasm and formed many puncta, ranging from circular spots to irregular patches (Figure 7A, ii). This construct behaved similarly to full-length Nup188 (Figure 7A, iii).

**Figure 7.**
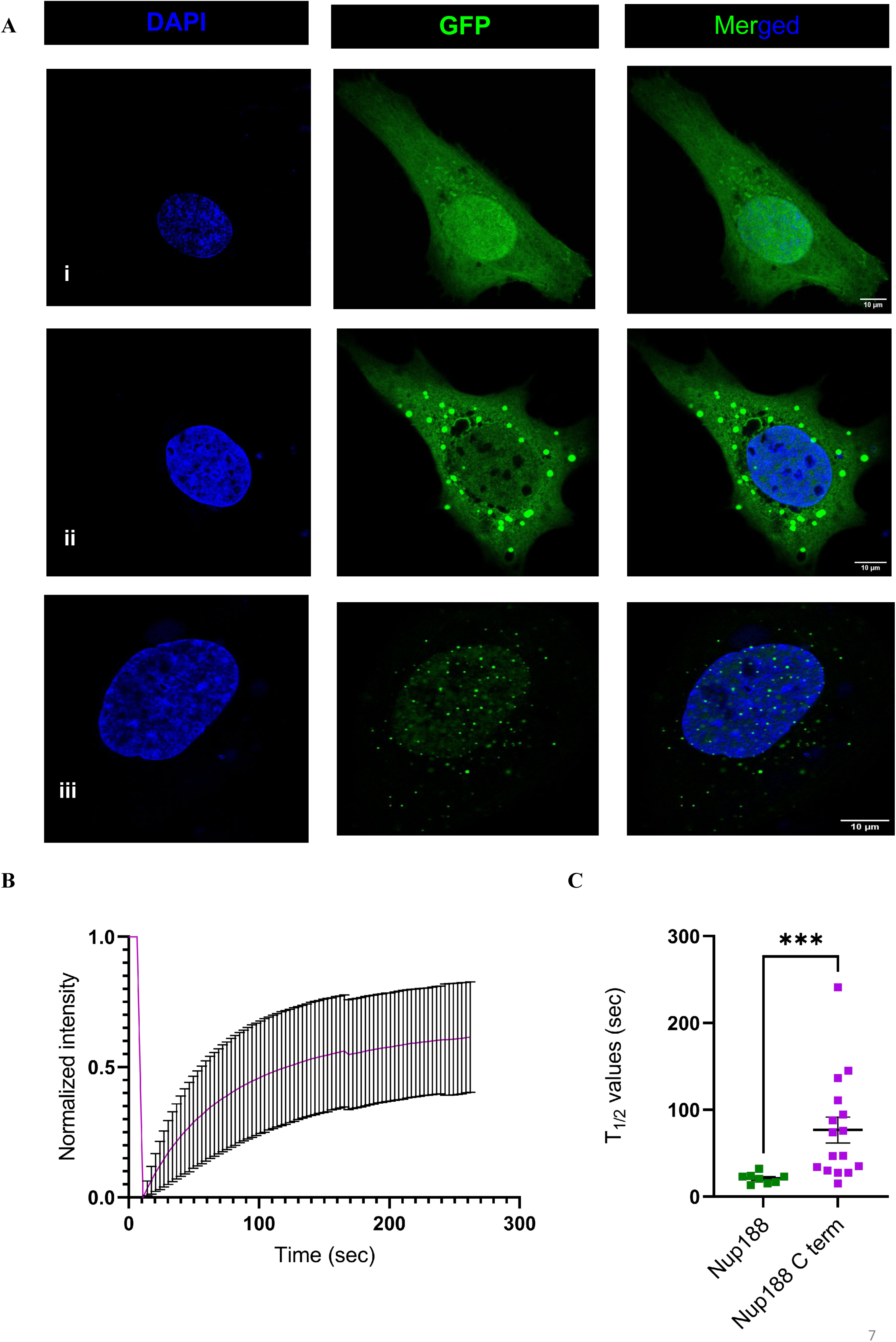
The C-terminal region of Nup188 drives condensate formation. (A) Confocal images of U2OS cells expressing GFP-Nup188 (1-1749) (i), GFP-Nup188 (1–1112) (ii) or GFP-Nup188 (1113–1749) (iii), stained with DAPI (blue). Only the C-terminal fragment forms condensates. Scale bars, 10 µm. (B) FRAP of GFP-Nup188 (1113–1749) condensates (mean ± SD, n = 17).(C) FRAP half-time (t½) of full-length Nup188 (n = 8) and C-terminal fragment (n = 17) condensates; Statistical comparison was performed using a two-tailed Mann-Whitney test; *** p = 0.0073, FRAP was performed one condensate per cell..

To further characterize the puncta formed by hNup188CTD-GFP overexpressed protein, we performed live cell imaging and FRAP experiments. We observed that the hNup188CTD-GFP fusion puncta are capable of moving, fusing, and dividing (Supplementary Figure S9B) and recovered after photobleaching, although more slowly than the full-length Nup188-GFP condensates (Figure 7B, 7C & Supplementary Figure S9C), suggesting the hNup188CTD-GFP condensates are far less dynamic than full length protein. Together this study reveals that the C-terminal domain of hNup188 is sufficient to nucleate condensate formation, however, it is less efficient than full-length hNup188 condensates.

### 8. MD simulations show that the three IDRs in the Nup188CTD contribute to condensate formation

As we observed that the hNup188CTD was sufficient to form condensates, we used multiscale molecular dynamics (MD) simulations to explore the underlying molecular basis. All-atom simulations were used to identify flexible regions, and coarse-grained simulations to test condensate formation. IUPred3 and FuzDrop predictions located the probable phase-separating regions mainly within residues 1372–1749 (Figure 1D, F). To capture this region while keeping microsecond-scale simulations computationally feasible, we simulated a shorter fragment, residues 1331–1749 (hereafter hNup188CTDshorter; Figure 8A), rather than the hNup188CTD (1113–1749), which was used in cell-based experiments. Initially, we generated the 3D model of hNup188CTD (UniProt ID: Q5SRE5) with AlphaFold3 (Abramson et al., 2024), which resulted in a model (pTM = 0.8) that appeared as a loop-rich extension appended to the folded α-solenoid core, with per-residue pLDDT scores falling locally below the confidence threshold (Figure 8A; Supplementary Figure S10).

**Figure 8.**
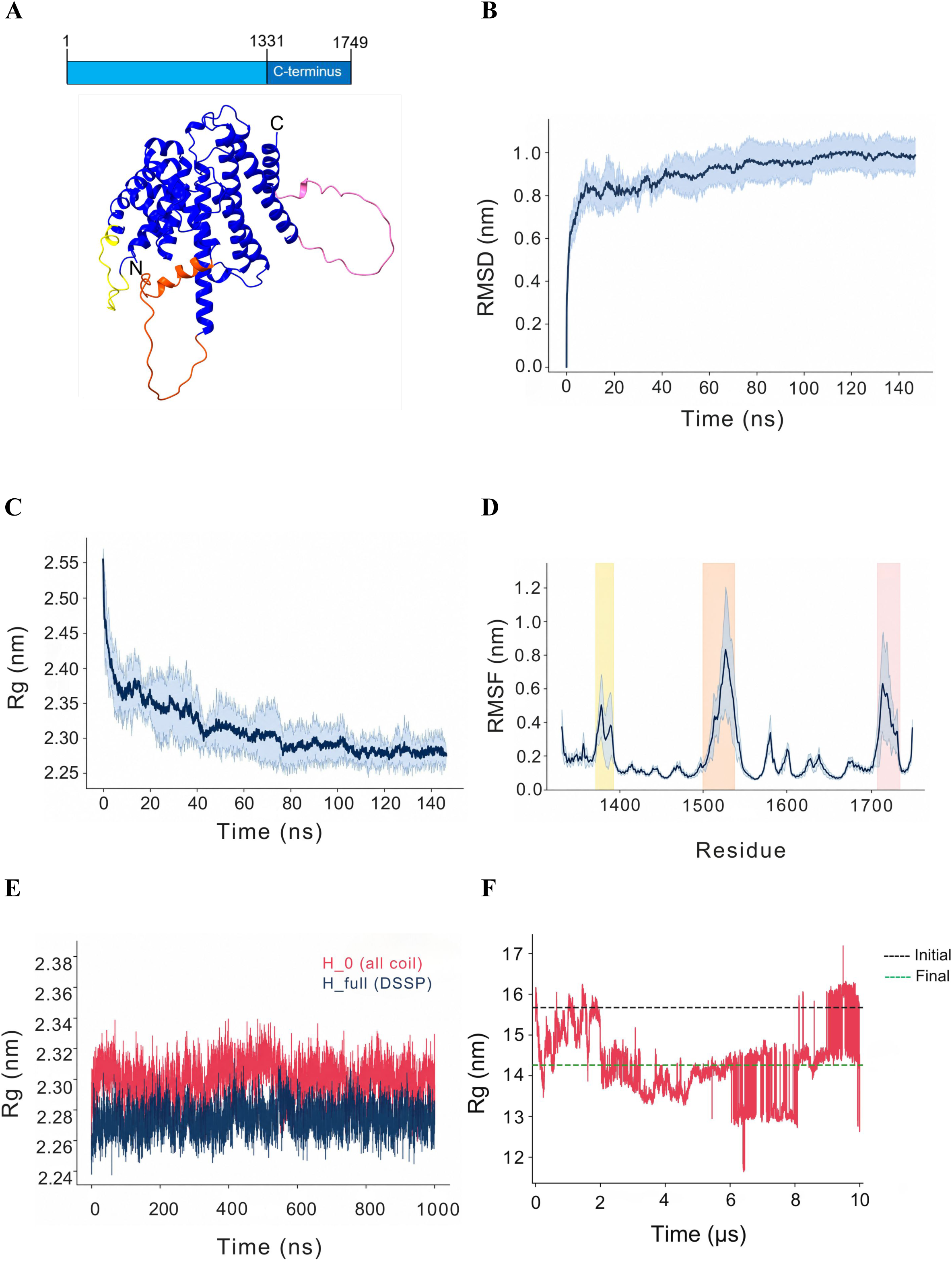
Molecular dynamics simulation of the Nup188 C-terminus region revealed three distinct IDRs. (A) Domain schematic of Nup188 with the C-terminal fragment (1331–1749) highlighted (top), and the AlphaFold 3 model of this fragment (pTM = 0.8) used for simulations (bottom). Helical regions are shown in dark blue. The three IDRs, defined by the RMSF peaks(Fig 1D), are coloured yellow (IDR1, 1371–1389), orange (IDR2, 1495–1535) and pink (IDR3, 1706–1734) respectively. (B–D) All-atom simulations (CHARMM36m; six independent 150-ns replicas). Mean ± 1 SD is shown for (B) Backbone RMSD, which plateaus at ∼0.95 nm after ∼70 ns, (C) Radius of gyration (Rg), which converges to ∼2.28 nm, and (D) Per-residue RMSF, showing ∼0.2 nm baseline flexibility with peaks at IDR1–3 (shaded, coloured as in A). (E) Single-chain Rg over 1 µs of Martini 3 coarse-grained simulation for H_0 (all coil, red) and H_full (DSSP-derived helices retained, blue). (F) Rg of the 24-mer C-terminus assembly over 10 µs of Martini 3 simulation, decreasing from 15.7 nm (black dashed line) to 14.25 nm (green dashed line).

To test whether these predicted low-confidence regions reflect genuine conformational flexibility, we performed all-atom simulations of the hNup188CTDshorter (six independent 150-ns replicas, CHARMM36m force field; see Methodology section). The backbone RMSD rose rapidly and plateaued at ∼0.95 nm by ∼70 ns (Figure 8B), and the radius of gyration (Rg) decreased from 2.55 nm to a converged value of ∼2.30 nm (Figure 8C), indicating that the fragment relaxed from its extended predicted conformation into a more compact, equilibrated ensemble. Per-residue root-mean-square fluctuation (RMSF) analysis showed a baseline flexibility of ∼0.2 nm across most of the chain, interrupted by three sharp, reproducible peaks, which we designate IDR1 (residues 1371–1389), IDR2 (1495–1535) and IDR3 (1706–1734) (Figure 8D). These peaks mark discrete sub-regions of elevated local mobility within the C terminus.

Secondary-structure propensity, intramolecular hydrogen bonding and solvent-accessible surface area across the six replicas (Supplementary Figure S11A–C) described a stable ensemble that interconverted between transient helical and coil states and plateaued by ∼80–100 ns. Intrachain contact maps showed dense short-range contacts along the diagonal and sparse long-range contacts (Supplementary Figure S11D). Together, these results indicate that the hNup188CTDshorter is compact but highly dynamic, particularly in the transition of few helical and coil states.

To confirm this behavior is retained in a coarse-grained analysis, which is better suited for larger-scale assembly simulations, we ran single-chain Martini 3 simulations (Souza et al., 2021) of the fragment, starting either from an all-coil conformation (H_0) or from a conformation in which the DSSP-derived helices from the all-atom trajectories were preserved (H_full). Both models converged to similar single-chain Rg values over 1 μs (Figure 8E; Supplementary Figure S12B), and both reproduced the similar regions (referred as IDR 1 – 3) of elevated flexibility (Supplementary Figure S12A). Rg of the 24-chain decreased from 15.7 nm to 14.25 nm over the simulation period (Figure 8F), indicating self-assembly. Thus, the coarse-grained model is consistent with the atomistic ensemble.

Together, these simulations support the view that the Nup188CTDshorter is a highly flexible, disorder-like segment, punctuated by three IDRs. This is consistent with its role as the condensate-driving domain in cells (Figure 7).

### 9. Coarse-grained simulations show the hNup188CTDshorter can phase separate

Because IDR containing sequences commonly drive biomolecular condensation (Banani et al., 2017.; Borcherds et al., 2021; Li et al., 2012), we asked whether the hNup188CTDshorter can assemble into a condensed phase *in silico* to further gain insights into the phase separation observed experimentally. We used coarse-grained simulations with two different force fields. In multichain Martini3 simulations of 24 copies of the fragment, the initially dispersed chains assembled into a single dense cluster over 10 μs (Figure 9A), consistent with progressive compaction and assembled into a condensate. The pairwise intermolecular contact map of the condensed phase revealed discrete residue clusters with markedly elevated inter-chain contact probability on a broadly contact-competent background. We observed that although most of the sequence can participate in inter-chain contacts, discrete “sticker” regions (IDR1–3) contribute disproportionately to assembly, consisting of the IDRs and surrounding helical regions (Figure 9B). In agreement, single-chain intrachain contact maps from the coarse-grained trajectories (Supplementary Figure S13A, S13B) showed a repeating, checkerboard-like pattern of long-range self-contacts in both the H_0 and H_full models, suggesting a sequence-encoded periodicity of self-association that is independent of residual secondary structure.

**Figure 9.**
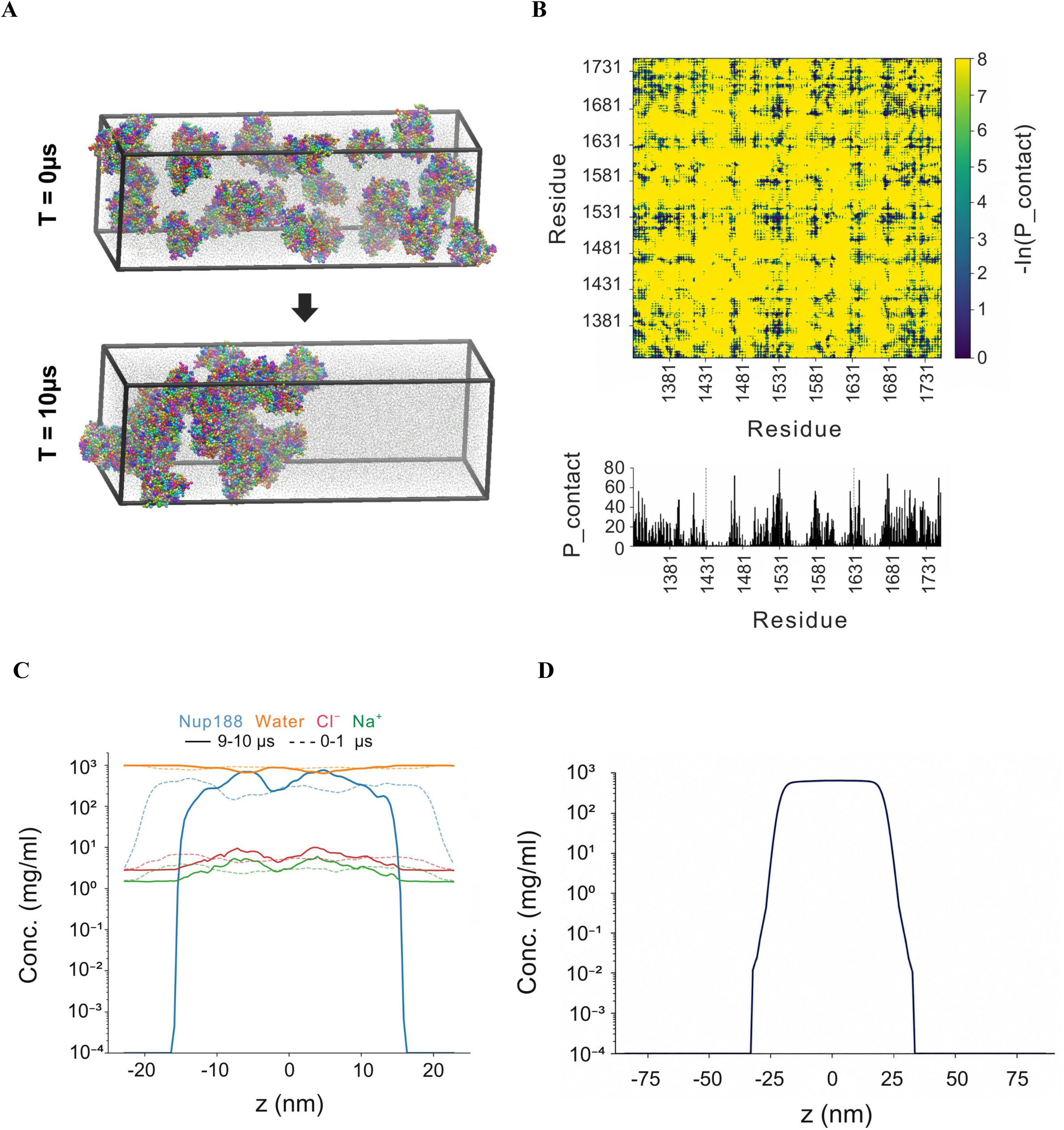
The Nup188 C-terminus undergoes phase separation in simulations. (A-C) Martini 3 coarse-grained simulations of 24 chains of the Nup188 C-terminal fragment.(A) Snapshots at 0 µs (top) and 10 µs (bottom), showing assembly into a single dense cluster. (B) Intermolecular contact map (top) and per-residue contact probability (bottom) in the condensed phase, plotted as −ln(P_contact); dark blue represents frequent contacts and yellow represents no contacts. (C) Density profiles along the slab z-axis (relative to the box centre) for Nup188, water, Cl⁻ and Na⁺, averaged over the first microsecond (dotted) and the last (9–10 µs, solid). (D) Nup188 concentration profile along z from HPS-Urry slab simulation.

To distinguish liquid-liquid phase separation from simple aggregation, we performed slab-geometry simulations also and computed density profiles along the long (z) axis of the simulation box. In the Martini3 slab simulations, the hNup188 concentration profile showed a plateau of high local concentration (Mean ∼454 mg/mL) at the center of the box, flanked by a dilute region at background concentration, whereas water and ion densities remained comparatively uniform (Figure 9C). This coexistence of a dense and a dilute phase is the defining signature of phase separation. To test whether the result depends on the force field, we repeated the slab simulation with the sequence-based HPS-Urry model (Regy et al., 2021) which likewise produced a well-defined high-concentration plateau (∼572±4 mg/mL) sharply demarcated from a dilute phase (Figure 9D). The two coarse-grained frameworks therefore agree that the hNup188 CTD has the propensity to phase separate. They differed, however, in the predicted molecular crosstalk. The inter-chain contact map from the HPS-Urry slab simulation (Supplementary Figure S14) showed contacts distributed diffusely and broadly across the sequence rather than concentrated at the discrete hotspots seen with Martini3. Martini3 thus favors a “stickers-and-spacers” organization, whereas HPS-Urry favors a more delocalized, additive interaction network. Overall, the simulations clearly established that the hNup188CTDshorter is capable of phase separation and IDR1-3 along with other transient interactions seem to drive this phenomenon.

### 10. Nup93 binds the Nup188 C-terminal region and suppresses Nup188 condensates

Nup93 is an established binding partner of Nup188 across species (Theerthagiri et al., 2010; Zabel et al., 1996). A recent cryo-electron tomography map of the human NPC (EMD-14322) captured the static interaction landscape of Nup188–Nup93 at low resolution and shows two copies of the Nup188–Nup93 in a single spoke (Figure 10A). To delineate the interaction interface, we performed pull-down assays with GFP-tagged truncation constructs of hNup188 (Supplementary Figure S15A) expressed in HEK293F cells and the GST-tagged anti-GFP nanobody. This analysis revealed that the full-length Nup188 interacted strongly with endogenous Nup93. Notably, among the fragments, we observed a faint band in the pulldown of endogenous Nup93 with the middle region (471–940) and the C-terminal region (1381–1749) (Figure 10B).

**Figure 10.**
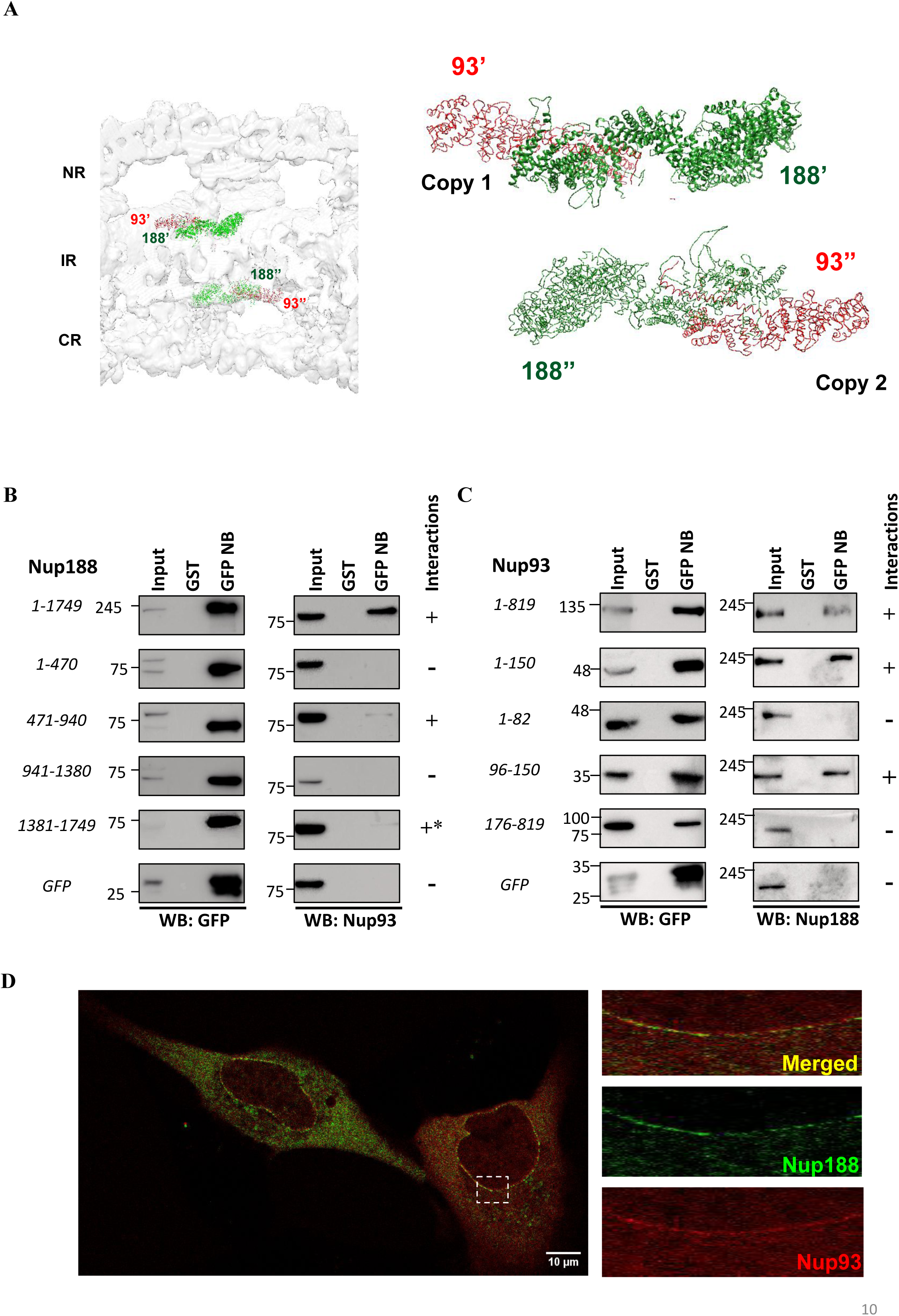
Mapping the Nup188–Nup93 interaction interface. (A) Placement of Nup188–Nup93 (two copies each) in the human NPC cryo-ET map (EMD-14322; NR, nuclear ring; IR, inner ring; CR, cytoplasmic ring), with the fitted models enlarged on the right. (B) GFP-nanobody (GFP NB) pull-down of full length (1–1749), 471–940 and 1381–1749 interact (shown with an asterisk (*)), whereas 1–470, 941–1380 and GFP alone do not.(C) Reverse pull-down with GFP-Nup93 fragments from cell lysates, probed for endogenous Nup188. Nup188 binds Nup93 1–819, 1–150 and 96–150, but not 1–82 or 176–819, mapping the interaction to residues 96–150.GFP-Nup188 fragments probed for endogenous Nup93; In B and C, GST beads and GFP alone are controls; +/− indicates interaction; molecular weights in kDa; n = 3 biological replicates. (D) U2OS cells co-expressing Nup93-mRuby (red) and Nup188-GFP (green). Nup188 no longer forms condensates and colocalizes with Nup93 at the nuclear envelope. The dashed box marks the enlarged region (merged and single channels). Scale bar, 10 µm. Representative of three independent experiments (10 images each).

In the reverse pulldown experiments, GFP-tagged fragments of human Nup93 (full-length, R1 [1–82], R2 [96–150], 1–150 and 176–819) (Supplementary Figure S15B) were expressed in HEK293F cells and pulled down to capture endogenous hNup188. We observed that the full-length hNup93 and the N-terminal 1–.150 fragment both interacted with endogenous hNup188, furthermore, we identified minimal hNup93 interacting region to 96–150 fragment, as it interacted efficiently with endogenous hNup188, whereas 1–82 and 176–819 couldn’t pull down endogenous hNup188 (Figure 10C). Based on this analysis, we concluded that the N-terminal 150 residues of Nup93, and in particular the R2 region (96–150) is responsible for binding to hNup188.

Based on our observation that hNup188 condensation is driven by its C-terminal domain (Figure 7) and hNup93 is likely to bind within this domain, we asked how hNup93 binding would affect the hNup188 condensates. When full-length hNup188–GFP was co-expressed with full-length hNup93-mRuby in U2OS cells, we observed that hNup188 no longer formed condensates and hNup93 also exhibited largely colocalized position with hNup188, (Figure 10D). In some cells, hNup188-GFP was also visible at the nuclear envelope, where hNup93-mRuby was also present. In control cells expressing Nup93-mRuby alone, Nup93 showed a diffuse cytoplasmic distribution with some signal at the nuclear envelope (Supplementary Figure S15C) and we observed no puncta formation. This suggests that possibly Nup93 binding to the Nup188 competes with Nup188 self-association and blocks condensate formation (Supplementary Figure S16).

## Discussion

Biomolecular condensates are membraneless structures, driven by multivalent, low-affinity interactions between IDR-containing proteins, that play a role in several biological processes. Among the ∼34 known Nups, about one third of them have IDR regions (FG repeats) (Beck & Hurt, 2017; Knockenhauer & Schwartz, 2016; Petrovic et al., 2022) and known to phase-separate (Dekker et al., 2023). Notably, hNup188 does not fall in this category, and our study revealed that even a non-FG-repeat-containing Nup is capable of forming condensate. Here we show that hNup188, a scaffold nucleoporin of the NPC inner ring, has the capacity to undergo liquid-liquid phase separation (LLPS) both *in vitro* and *in cellulo* (Figure 2; Supplementary Figures S3 & S4). The Nup188 condensates showed similar material properties like FXFG Nup - Nup98 (Figure 4 and 5). Sequence-based predictors flagged a disordered C-terminal of Nup188 as condensate-promoting (Figure 1D–1F; Supplementary Figure S2). Furthermore, using MD simulations we identified flexible regions and sticker-like segments (IDR1–3) within it and demonstrated how it is contributing to phase separation (Figures 8 and 9). We also report that co-expression of hNup93 (Nup93 itself does not exhibit condensate formation) suppressed condensate formation when it is interacting to this region (Figure 10), indicating a critical role of the Nup188-Nup93 interface in regulating the LLPS behaviour of hNup188.

### A structured scaffold Nup with FG-Nup-like condensate properties

Scaffold nucleoporins are built from stacked α-helical solenoids and β-propellers and form the symmetric core of the pore (Fernandez-Martinez & Rout, 2021). In contrast, the intrinsically disordered FG repeat-containing Nups are known to form liquid droplets and hydrogels that reproduce the selectivity of the NPC permeability barrier (Celetti et al., 2020; Frey et al., 2006; Schmidt & Görlich, 2015). Condensation of FG Nups has also been reported to be linked to NPC biogenesis in a few organisms (Hampoelz et al., 2016, 2019; Lin et al., 2026). In the case of yeast, it is reported that Nsp1 foci/condensates rarely contain the scaffold nucleoporins such as Nup133 and Nic96 (Otto et al., 2024). To our knowledge, condensate formation by an inner-ring scaffold nucleoporin on its own has not been reported yet. Our data on hNup188 suggest that the boundary between “scaffold” and “condensate-forming” nucleoporins is far less sharp than assumed. The FRAP half-times of Nup188 and Nup98 condensates were observed to be very similar (median 21.80 s versus 22.66 s), and FlickerPrint analysis placed most condensates of both proteins in a low-interfacial-tension, low-bending-rigidity regime with near-circular shapes (Figures 4F & 5; Supplementary Figures S7 and S8). A previously reported study on fungal Nup188 and its paralog Nup192 (Nup205 in vertebrates) showed that both are stacked helical proteins with intrinsic flexibility that resemble nuclear transport receptors (NTRs), and can cross the pore by interacting with FG Nups via facilitated diffusion (Andersen et al., 2013). Thus, it can be speculated that hNup188 may interact directly with FG-nucleoporin condensates, and it is a small step from interacting with condensates to forming them.

### Parallels with other helical proteins that phase separate

Our study presented here places Nup188 alongside a growing set of proteins in which helical elements contribute to biomolecular condensate formation. The best characterized example is TDP-43, whose disordered C-terminal fragment contains a conserved partially α-helical region. Helix–helix contacts within this region drive liquid–liquid phase separation. Recent structural work indicates that hydrophobic residues drive this helix–helix assembly, while polar residues dampen it (Rizuan et al., 2022). The Nup188 C-terminus resembles this situation in several respects. In our all-atom simulations, it was neither a stable fold nor a random coil: it interconverted between transient helical and coil states, contained three sharply flexible sub-regions (IDR1–3), and the coarse-grained models placed elevated inter-chain contact probability in discrete sticker-like segments (Figures 8 and 9; Supplementary Figures S11 and S13), thus highlighting the common rules for condensate formation among similar proteins.

Secondly, it is known that extended coiled-coils can act as multivalent stickers with defined geometry. For example, the centrosome scaffold protein SPD-5, which contains nine predicted coiled-coil domains, forms micron-scale condensates in vitro whose assembly depends on multivalent coiled-coil interactions and is regulated by PLK-1 phosphorylation (Rios et al., 2024; Woodruff et al., 2017). Although hNup188 does not contain a coiled-coil motif, however, the principle carries over: structured helical elements can provide directional, regulatable multivalency that a purely disordered chain cannot provide. Interestingly, both the centrosome and the NPC are large proteinaceous scaffolds built from many helical proteins, and in both, condensation may serve as a route to assembly that is regulated by binding partners or post-translational modifications. Notably, Nup188 has been reported to form barrel-like structures around centrioles at the ciliary base (del Viso et al., 2016), and localizes at the centrosome (Itoh et al., 2013), plausibly performing these moonlighting functions based on liquid-liquid phase separation interactions.

### A cytoplasmic pool distinct from annulate lamellae

We show that the endogenous hNup188 localized to the nuclear envelope in WI-38 fibroblasts, but in HeLa and U2OS cells it was present as mainly punctate, localized in the nucleus and cytoplasm respectively, with no evident nuclear envelope staining in U2OS cells (Figure 3; Supplementary Figure S5). The distinct spatial localization of the resulting nuclear puncta in HeLa and cytoplasmic in U2OS - indicates cell-line-specific differences in the local microenvironment. Altogether, these observations point to an NPC-independent pool of Nup188. Cytoplasmic nucleoporin assemblies often annulate lamellae, and Nup358-containing condensates are precursors of annulate lamellae as shown in oocytes (Hampoelz et al., 2016, 2019). A large subset of Nups has been already shown to be part of these ALs (Lin et al., 2026) including the hNup188 paralog Nup205. Consistent with its role as a bona fide AL component, hNup205 localized to ALs in our study also; however, Nup188 condensates failed to colocalize with the AL marker Nup358 (Figure 6). The two paralogs therefore appear to have different non-NPC destinations. The function of the Nup188 condensate pool is not known yet. However, it is reported that hNup188 is a component of condensate assemblies such as centrosomes and cilia (del Viso et al., 2016; Kostiuk & Khokha, 2021; Vishnoi et al., 2020; Woodruff et al., 2017; Zwicker et al., 2014). Rather than serving merely as a storage reservoir for scaffold nucleoporin supply during pore biogenesis, these hNup188 condensates may carry out specialized, non-NPC functions directly at centrosomes or basal bodies.

Since hNup188 has multiple roles inside the cell, pathological shifts in the level of hNup188 expression or phase separation may directly drive cellular dysfunction and disease pathogenesis. Indeed, hNup188 overexpression is reported across multiple malignancies (Zhu et al., 2025), whereas truncated protein leads to loss of function causing Sandestig-Stefanova syndrome, a severe developmental disorder (Muir et al., 2020; Sandestig et al., 2020; Almalki et al., 2025 and Soğukpınar et al., 2026). However, how hNup188 dependent processes drive these clinical phenotypes and how its liquid-like condensate behavior contributes to their pathologies remains unanswered.

### hNup93 as a disrupter of hNup188 condensation

Our data presented here revealed that the R2 region of hNup93 is the minimal interacting region for hNup188 binding (Figure 10B, 10C). This agrees with previous structural studies in which an extended N-terminal helix of Nup93 inserts into a groove of Nup188 (Lin et al., 2017; Petrovic et al., 2022). Notably, we also observed that full-length hNup188 binds Nup93 more efficiently than its CTD alone (Figure 10C). The hNup188CTD includes IDR1–3 regions, which significantly contribute to transient interactions for condensate formation (Figure 9). Interestingly, co-expression with hNup93 abolished hNup188 condensate assembly. Based on this, we propose a model that the hNup93 R2 region occupies the C-terminal region and competes with the homotypic contacts of Nup188CTD-Nup188CTD that drive condensation (Supplementary Figure S16).

Regulation of condensation by helical binding partners has been known in the field. For example, the Nuclear Transport Receptors (NTRs), which are helical solenoids, bind their cargo at specific sites and additional weaker sites, and thereby prevent and reverse the phase separation of FUS and other RNA-binding proteins (Guo et al., 2018; Hofweber et al., 2018; Qamar et al., 2018). Additionally, in oocytes, Ran and NTRs also regulate nucleoporin condensates during NPC biogenesis (Hampoelz et al., 2019). Given the structural similarity between Nup188 and NTRs (Andersen et al., 2013; Flemming et al., 2012), Nup93 may play a comparable, chaperone like role for Nup188, holding it soluble until it is incorporated into the pore.

### Potential liquid like proteins as Nup188 interactors

To put our findings in a wider context, we predicted the LLPS propensity of hNup188 interactors listed in BioGRID (thebiogrid.org) using MolPhase. Out of 205 reported interactors, 138 proteins showed the predicted score exceeding 0.5 (Supplementary Figure S17). This suggests that Nup188 may be a part of a wider network of proteins with condensate-forming potential. It should be noted that these interactors are compiled from a variety of cellular conditions (Go et al, 2021; Lee et al., 2021; Cho et al., 2022; Siebert et al., 2022), so direct experimental validation of these candidate interactions, and testing whether any of them modulate Nup188 condensate formation in the manner we describe for Nup93, represents an important direction for future work.

### Limitations of the study

Although we clearly establish the liquid-like property of the hNup188 protein, there are yet many questions to be answered. Firstly, the specific role of the identified Nup188 IDR1–3 and other helical regions in condensate formation and their cellular relevance is yet to be elucidated. Moreover, whether FG/GLFG Nups condensate interact with hNup188 needs to be investigated. On top of that, the extent of non-NPC-related functions of hNup188 driven by condensate physiology needs to be addressed in the future so that its role in human diseases could be fully documented. Also, the suppression of condensation by Nup93 was shown by co-expression, the underlying mechanism, competition versus altered localization, remains to be directly established.

## Methodology

### 1. Computational analysis of hNup188 sequence

All computational analysis - IUPred3, FuzDrop and MolPhase (See Supplementary Table) were carried out by using UNIPROT protein inputs as: human Nup188 protein (ID: Q5SRE5), human Nup98 (ID: P52948) and human Nup205 (ID: Q92621). The output obtained in csv format from each tool has been used for plotting graphs using GraphPad Prism 9.

### 2. Cloning, Expression and Purification of hNup188

Full length human Nup188 (residues 1-1749) was cloned into a pEG BacMam vector containing GFP and 8X His tag. The gene was followed by a TEV protease cleavage site. The details of primers and restriction sites used for the cloning are given in the Supplementary table. HEK293F suspension cells were grown in FreeStyle 293 media (Gibco) at 37 °C with 8 % CO_2_ and 80 % humidity with a shaking of 110 rpm. For large scale expression of the protein, pEG BacMam Nup188 FL plasmid was transfected into the HEK293F cells of density 1.0 to 1.5 million using PEI in 1:3 ratio. Post 72 hrs, the cells were harvested and resuspended in a resuspension buffer (50 mM Tris Cl – pH 7.5, 300 mM NaCl, 2 % DDM, 5 mM β-Me, 1 mM EDTA, 1x PIC, 1 mM MgCl_2_ and 10 % glycerol). Following resuspension, the cells were lysed using sonication (35 amp, 10 s on 20 s off) and clarified further by high-speed centrifugation at 4 °C. The obtained supernatant was bound to GST beads which are already bound with GST-tagged anti-GFP nanobodies for 2 hours at 4 °C in an end-to-end rotor. The beads were washed with 100 CV of washing buffer (50 mM Tris Cl - pH7.5, 300 mM NaCl, 0.5 % DDM, 5 mM β-Me, 1mM EDTA, 1 mM MgCl_2_ and 5% glycerol). The bound protein was eluted in elution buffer (50 mM Tris Cl - pH 7.5, 300 mM NaCl, 5 mM β-Me, 1 mM DDM, 1mM EDTA, 1 mM MgCl_2_, 5% glycerol, 10mM reduced glutathione). The affinity purified protein was subjected to PreScission protease treatment to remove the GST tag. The protein thus obtained was further purified using gel filtration chromatography using Superose 6 increase 10/300 24 mL column, which is already equilibrated with SEC buffer (50mM Tris Cl - pH 7.5, 300 mM NaCl, 0.17 mM DDM, 1 mM EDTA, 2 % Glycerol, 1 mM MgCl_2_, 1mM DTT). The eluted fractions were run on 10 % SDS PAGE to confirm the molecular size of the protein.

### 3. In vitro condensation assays

Recombinantly expressed, size-exclusion chromatography (SEC)-purified GFP-tagged hNup188 was prepared at a working concentration of 0.2 mg/mL (∼1 µM). Reaction mixtures with a final volume of 20 µL were assembled at 4 °C across a series of PEG-5000 crowding concentrations (0%, 1%, 2.5%, 5%, and 10% w/v). To negate possible effects of the detergent in condensate formation, additionally GFP alone was also assayed in an identical DDM-containing buffer.

### 4. Cell culture

HEK293FT, HeLa and U2OS cells were cultured in Dulbecco’s modified Eagle’s medium (DMEM) supplemented with 10 % fetal bovine serum (FBS), with Penstrep at 37 °C in 5 % CO_2._ The cells are passaged every 2 - 3 days after dissociation using 1X TrypLee in 1:4 ratio. Human embryonic lung fibroblasts (WI-38) were maintained in Minimum Essential Medium (MEM) supplemented with complete growth factors, including 15% FBS, 1x non-essential amino acids, and 1x anti-anti, and sub cultured when the desired confluency was reached. For passaging, the culture medium was aspirated, and the cell monolayers were incubated with 0.25% Trypsin-EDTA for 3 minutes at 37 ⁰C to facilitate detachment. The enzymatic activity was subsequently neutralized by adding an equal volume of fresh, complete MEM. The dissociated cell suspension was collected into a conical centrifuge tube and pelleted by centrifugation at 2000 RPM for 3 minutes. The resulting cell pellet was resuspended in fresh complete MEM and split at a ratio between 1:3 and 1:4.

### 5. Immunofluorescence assay

U2OS/WI-38/HeLa human lung fibroblasts were cultured and harvested at approximately 60 − 70% confluency. The growth medium was aspirated, and the cells were gently rinsed with 1× phosphate-buffered saline (PBS). Fixation was performed by incubating the cells with 4% paraformaldehyde (PFA) in PBS for 15 minutes at room temperature under continuous agitation on a laboratory shaker. Following fixation, the PFA solution was removed, and the cells were subjected to three consecutive washes with 1× PBS for 5 minutes each. To facilitate intracellular detection, cells were permeabilised by incubation with 0. 5% Triton X-100 in 1× PBS for 12 minutes on ice, followed by three additional 5 min washes with 1× PBS. Non-specific binding sites were subsequently blocked by incubating the cells with a blocking buffer consisting of 3% fetal bovine serum (FBS) in PBS for 1 ℎ at room temperature. For target detection, cells were incubated overnight at 4^◦^*C* with a rabbit anti-NUP188 primary antibody (Cat#ab8644: dilution 1:150) diluted in blocking buffer to label endogenous NUP188. For membrane labelling, we used mab414 (1:400). Unbound primary antibodies were removed by washing the cells three times for 5 minutes each with 1× PBS contains 0.1% Tween-20 (PBST). Cells were then incubated with an Alexa Fluor anti-rabbit 488 (Cat no: A21206, Dilution-1:1000) secondary antibody (diluted in blocking buffer) for 45 minutes at room temperature, protected from light. Residual secondary antibody was removed via three 5-minute washes with 1× PBST. Nuclei were counterstained by incubating the cells with Hoechst dye diluted in PBS for 5 minutes, followed by a final washing cycle consisting of three 5-minute washes in 1× PBST. Finally, the cells were mounted onto glass slides using a suitable antifade mounting medium. High-resolution confocal images were acquired using an Olympus FV3000 Confocal Laser Scanning Microscope (CLSM) and subsequently processed and analysed utilising Olympus cellsens software. For the structured illumination microscopy (SIM) experiments, we have transiently transfected U2OS cells with vectors expressing Nup188 and Nup205 proteins and fixed and processed them as described above and imaged using Nikon V3-Deep SIM microscope. The images captured were reconstructed using NIS Elements imaging software version 6.20.00.

### 6. Fluorescence Recovery After Photobleaching (FRAP) and 1,6 hexanediol assay

U2OS cells were seeded on a glass coverslip, allowed to adhere for 16 hours, and then transfected with fluorescently-tagged Nup188 DNA using PEI at a 3:1 ratio. Twenty-four hours post-transfection, the cells underwent FRAP: three initial frames were collected as the pre-bleach baseline, followed by a 1-second pulse of 100% laser power to photobleach the target focus. Fluorescence recovery was then recorded over an additional 40 post-bleaching frames, with the resulting recovery curve plotted in Excel to analyze the molecular mobility. In case of hexanediol experiment, following 36hrs of post transfection. U2OS cells were imaged for 10 frames to establish a pre-treatment baseline. 1,6-Hexanediol was then added to a final concentration of 5%, and imaging continued for an additional 10 frames to monitor condensate dissolution.

### 7. Flickerprint analysis

Condensate material properties and shape descriptors were quantified using Flickerprint, a Python package that applies flicker spectroscopy to fast time-lapse microscopy videos of condensates to extract bending rigidity (κ), interfacial tension (σ), and morphological parameters (circularity, solidity, axis ratio), as described previously (Williamson et al., 2025). Briefly, Live-cell videos of U2OS cells transiently expressing either Nup188-GFP or Nup98-GFP were acquired on a Spin SR spinning-disk confocal microscope, using a 60× objective, at the maximum frame rate achievable by the acquisition system. Videos were processed using Flickerprint following the filtering and analysis pipeline described in Williamson et al., 2025. Density scatter plots and frequency histograms were generated in Python using Matplotlib.

### 8. Molecular Dynamics simulations

MD simulations were performed by using Nup188 C-terminus (residues 1331–1749; pTM = 0.8), retrieved from the AlphaFold3 (Abramson et al.,2024) model of full-length Nup188 (UniProt ID: Q5SRE5). Nup188 dynamics were evaluated initially at the all-atom level. Systems were built with CHARMM-GUI (Jo et al.,2008) using the CHARMM36m (Huang et al.,2017) force field. The monomer was solvated in a cubic box of TIP3P water (Jorgensen et al.,1983) with a 1.2 nm buffer around the protein and neutralized with 150 mM KCl. Each system was energy-minimized by steepest descent until the maximum force fell below 1000 kJ mol⁻¹ nm⁻¹. It was then equilibrated for 100ps under NVT conditions (v-rescale thermostat (Bussi et al., 2007), 310K, τ_T = 0.1 ps) and 100 ps under NPT conditions (Berendsen barostat (Berendsen et al.,1984), 1 bar, τ_P = 2.0 ps). Heavy-atom position restraints (1000 kJ mol⁻¹ nm⁻²) were applied during both equilibration steps. Production simulations used the v-rescale thermostat and the Parrinello–Rahman barostat (Parrinello and Rahman, 1981) (τ_P = 5.0 ps; compressibility 4.5 × 10⁻⁵ bar⁻¹), with a 2-fs time step. Bonds involving hydrogen were constrained with LINCS (Hess et al.,1997). Six independent replicas, started with different initial velocities, were run for 150 ns each (900 ns in total) in GROMACS 2021.5 (Abraham et al., 2015).

Coarse-grained (CG) topologies were generated with martinize2 (Kroon et al.,2023) for the Martini 3.0.0 force field (Souza et al.,2021), with side chain corrections by scfix (Herzog et al., 2016). An elastic network (0.5–0.9 nm cutoffs; force constant 500 kJ mol⁻¹ nm⁻²; Periole et al.,2009) was applied to maintain the folded regions. Two secondary-structure variants were prepared: H_full, which retained the DSSP (Dictionary of protein secondary structure) assigned helices (Kabsch and Sander,1983) and H₀, in which all residues were modeled as coil. Systems were solvated with insane.py (Wassenaar et al.,2015) containing 150 mM NaCl. Simulations used the v-rescale thermostat at 310 K (Bussi et al., 2007), the Parrinello–Rahman barostat at 1 bar (Parrinello and Rahman, 1981), reaction-field electrostatics (new-rf parameters) (de Jong et al.,2016) and a 20-fs time step. Three systems were simulated, each in three replicas: a single chain (10 × 10 × 10 nm box, 1 µs) and 24 chains arranged 2 × 2 × 6 in an elongated box (16 × 16 × 48 nm, 10 µs). As HPS-Urry one-bead-per-residue model is extensively used for phase separation-related simulations (Regy et al., 2021; Dignon et al., 2018; Dignon, Best and Mittal,2020; Ingólfsson et al.,2023), we performed slab simulations using HOOMD-blue 2.9.7 (Anderson, Glaser and Glotzer,2020) (CUDA 10.2 GPU build). One hundred chains were placed in a 20 × 20 × 168 nm slab box with periodic boundary conditions. Folded regions were treated as rigid bodies (hoomd.md.constrain.rigid). Simulations were run at 310 K with a Langevin thermostat (τ = 1000 ps) and a 10-fs time step, for 3.5 µs.

All-atom and Coarse-grained simulations were analysed using GROMACS tools for standard analysis and custom analysis was done by using MDAnalysis (Michaud-Agrawal et al.,2011), NumPy (Harris et al.,2020) and Matplotlib (Hunter,2007). Inter-residue contacts were counted with a 0.8-nm distance cutoff. Secondary structure was assigned with DSSP (Kabsch and Sander,1983) algorithm and the snapshots were made with VMD (Humphrey, Dalke and Schulten,1997).

### 9. Pull-down assays

The pull-down assay was performed as described in (Chopra et al., 2020). Briefly, ∼1-1.5 mg of protein lysate was incubated with purified GST-tagged anti-GFP nanobody (construct in pGEX-6P1) (Addgene ID # 61838) bound glutathione beads. The proteins were probed with primary antibodies: anti-Nup188 (Abcam ab86601) at 1:4000 dilution, anti-Nup93 (Abcam Ab53750) at 1:500 dilution and anti-GFP (Sigma G1546) at 1:3000 dilution. The blots were developed using Quant HRP substrate (Takara) and images were acquired on Amersham Imager 600 (GE healthcare).

### 10. Phase separation propensity analysis for Nup188 protein in BIOGRID interaction network

Protein-protein interaction data for human Nup188 protein (Uniprot ID: Q5SRE5) was retrieved from the BioGRID database (BioGRID version 5.0.255) using Nup188 as the query protein. All reported interactions were extracted and their intrinsic propensity to form liquid-liquid phase separation (LLPS) was predicted using MolPhase prediction server (https://molphase.sbs.ntu.edu.sg/). Proteins with a MolPhase score above 0.5 (tool’s recommended cut-off) were classified as having LLPS propensity. The Nup188 interactome, along with their corresponding MolPhase-derived LLPS propensity scores, was visualized as a protein–protein interaction network using Cytoscape (version 3.10.4), where, Nup188 was set as the central node, connected to each of its identified interactors by an edge representing the corresponding BioGRID-reported interaction. Nodes were colored according to their MolPhase LLPS propensity scores to visually distinguish high- and low-propensity interactors within the network.

## Supplementary table

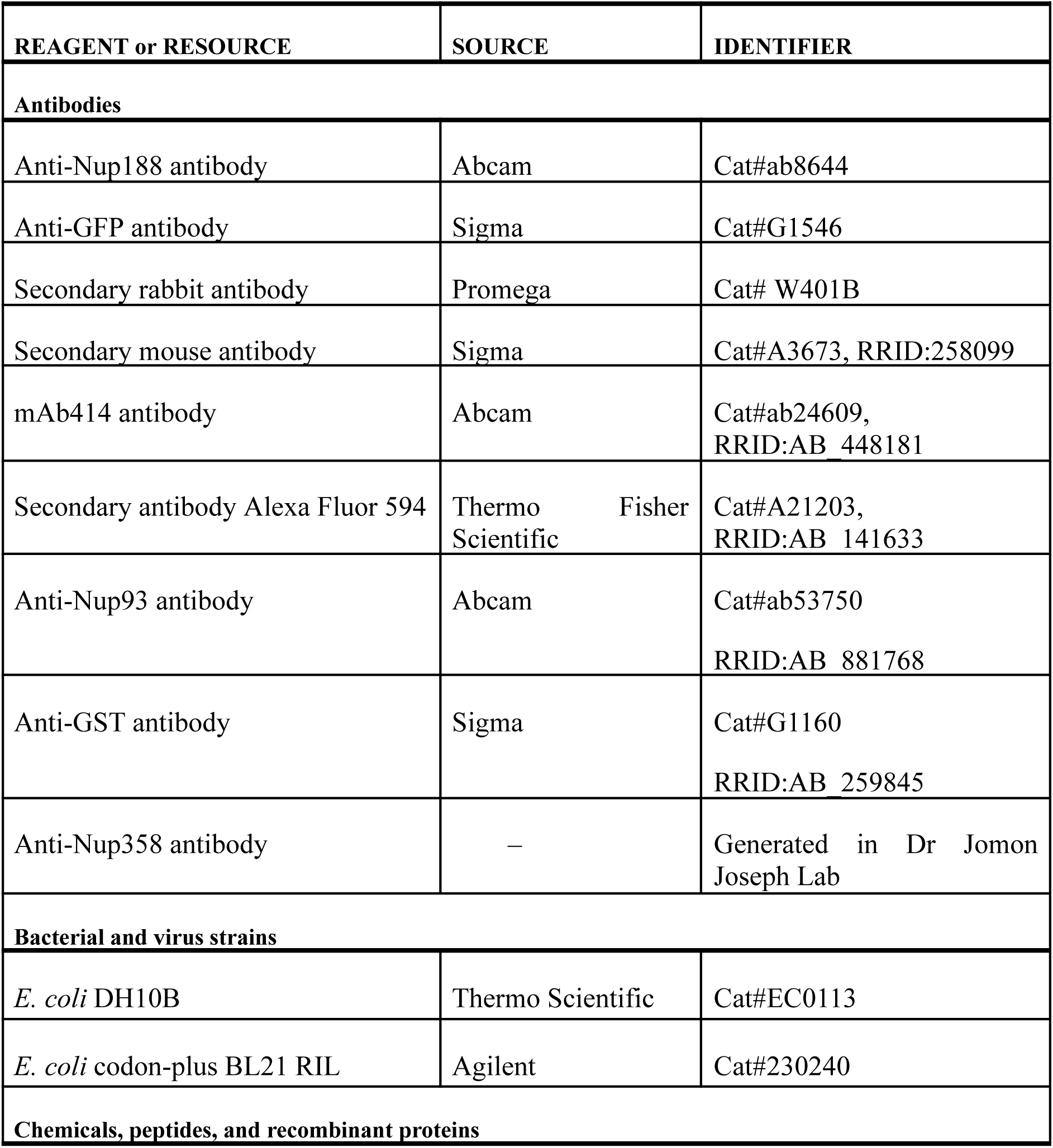
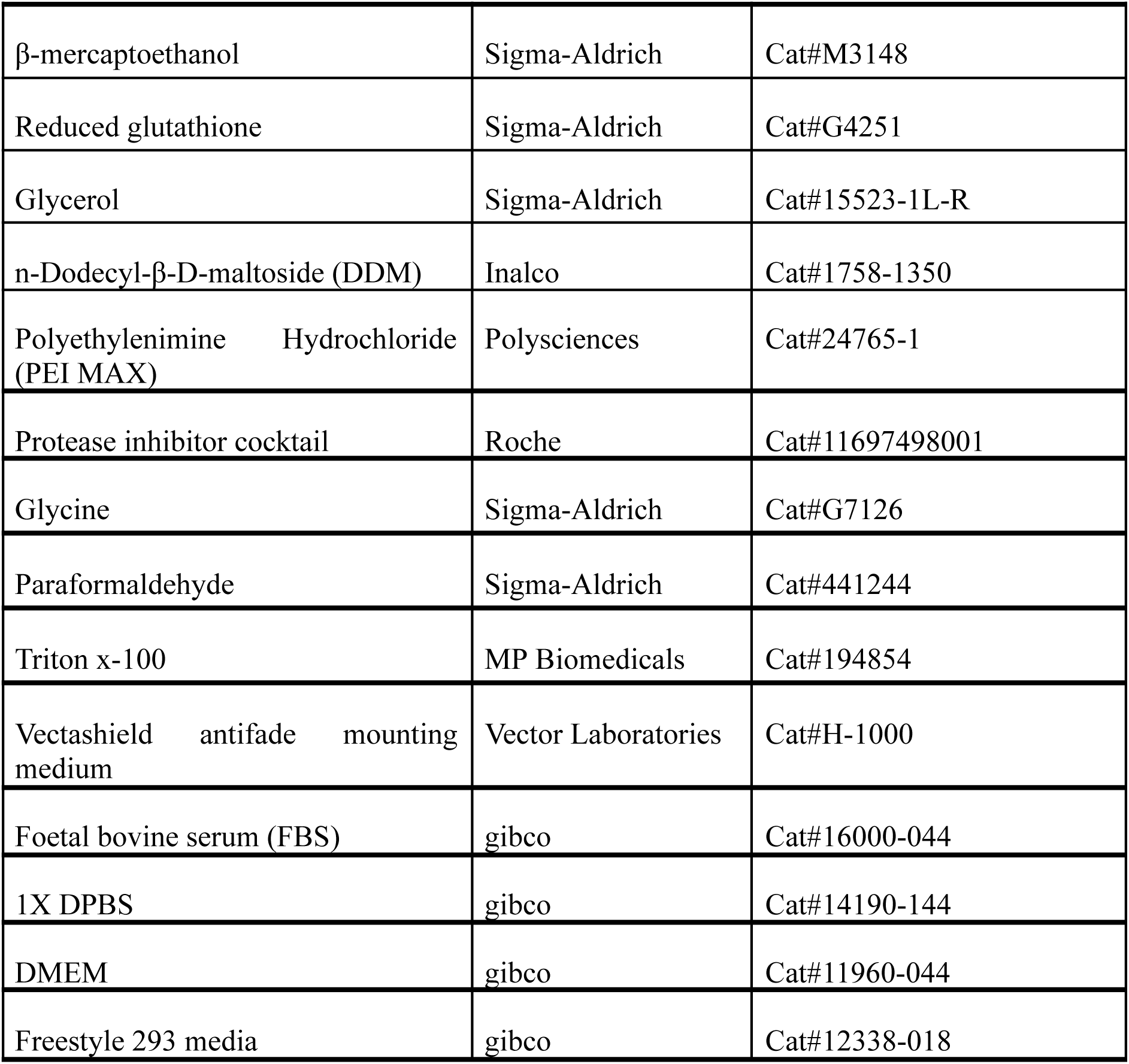

| Experimental models: cell lines |  |  |
| --- | --- | --- |
| Human: HEK293F cell line | Invitrogen | Cat#R79007 |
| Human: HEK293T cell line | ATCC | Cat#CRL-3216 |
| U-2 OS | ATCC | Cat#HTB-96 |
| HeLa | ATCC | Cat#CRM-CCL-2 |
| WI-38 | ATCC | Cat#CCL-75 |
| Recombinant DNA |  |  |
| pGEX-6P-1-GFP-nanobody | Addgene | Cat#61838 |
| Plasmid - hNup188 (1-1749) in pEG BacMam GFP His | This paper | N/A |
| Plasmid - pEGFP-C1 | Addgene | Cat#6084-1 |
| Plasmid - pEGFP-C1hNup98(1-937) | This paper | N/A |
| Plasmid - pEGFP-C1 hNup188(1-1749) | This paper | N/A |
| Plasmid - pEGFP-C1 hNup188(1113-1749) | This paper | N/A |
| Plasmid - pEGFP-C1 hNup188 (1-1112) | This paper | N/A |
| Plasmid- pEGFP-C1 Nup93 (1-819) | Chopra et al., 2020, Sonawane et al., 2020 | N/A |
| Plasmid- pEGFP-C1 Nup93 (1-150) | Chopra et al., 2020, Sonawane et al., 2020 | N/A |
| Plasmid- pEGFP-C1 Nup93 (1-82) | Chopra et al., 2020, Sonawane et al., 2020 | N/A |
| Plasmid- pEGFP-C1 Nup93 (96-150) | Chopra et al., 2020, Sonawane et al., 2020 | N/A |
| Plasmid- pEGFP-C1 Nup93 (176-819) | Chopra et al., 2020, Sonawane et al., 2020 | N/A |
| Plasmid -pEG BacMam mRuby StrepII - hNup93 (1-819) | This paper | N/A |

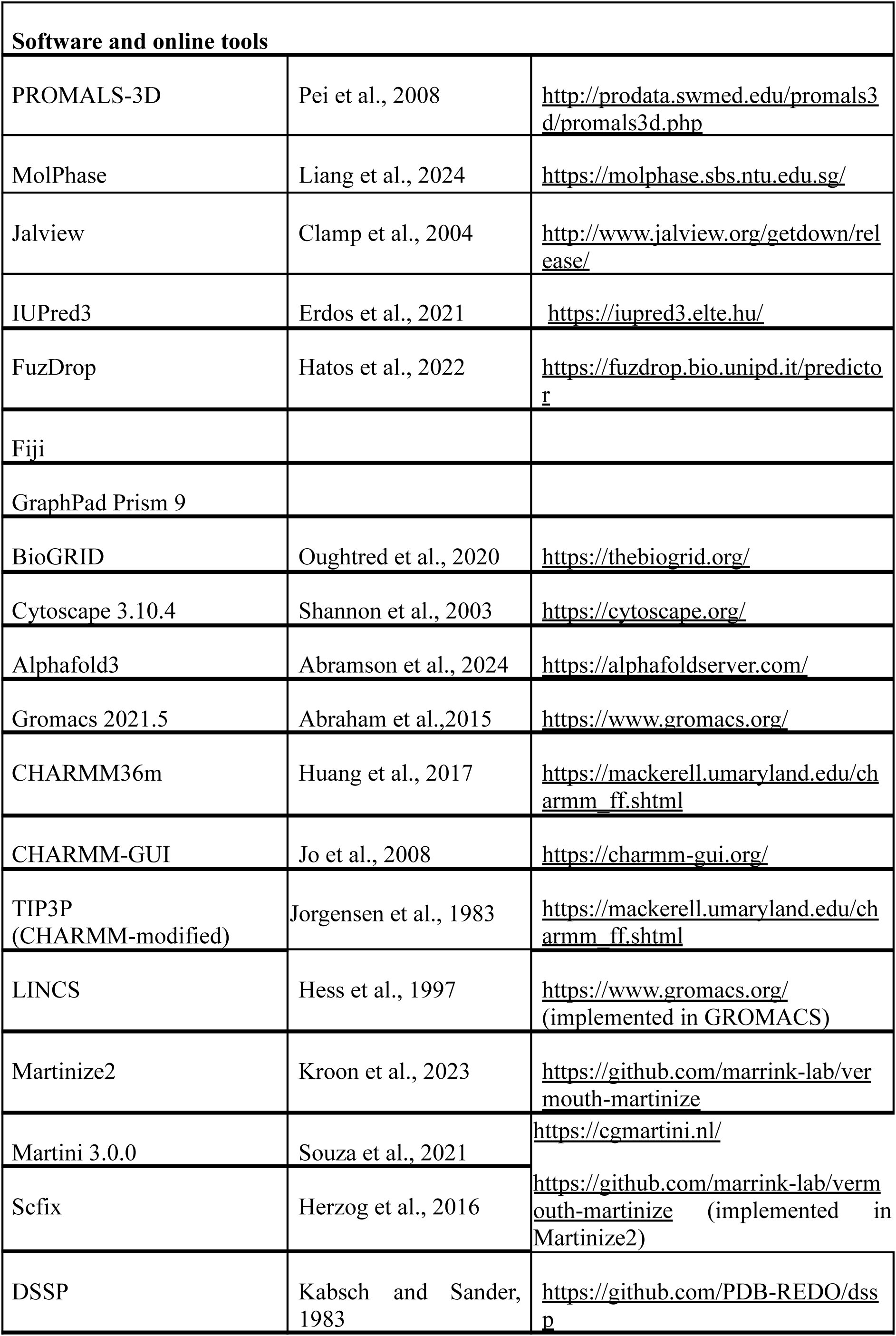
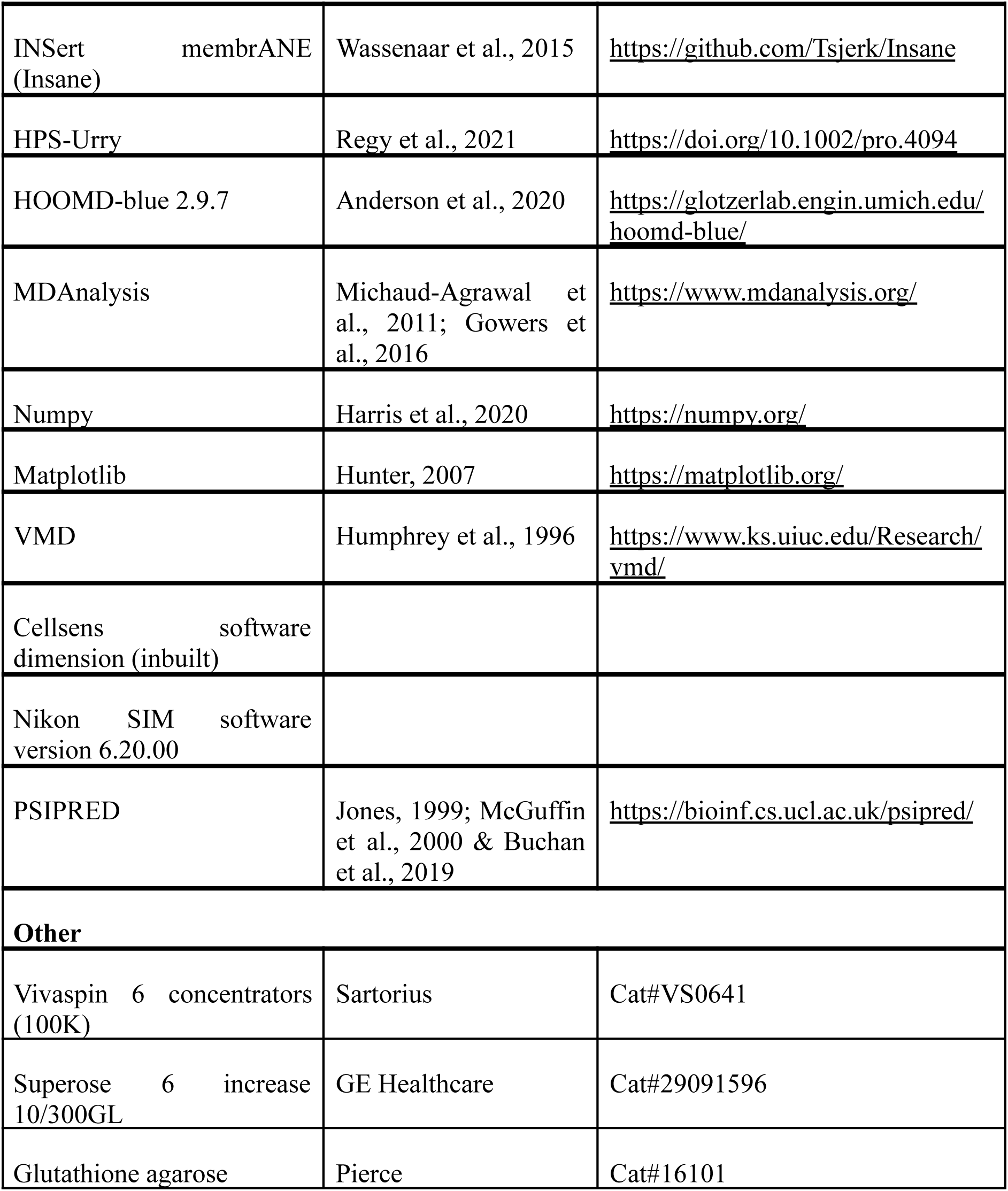

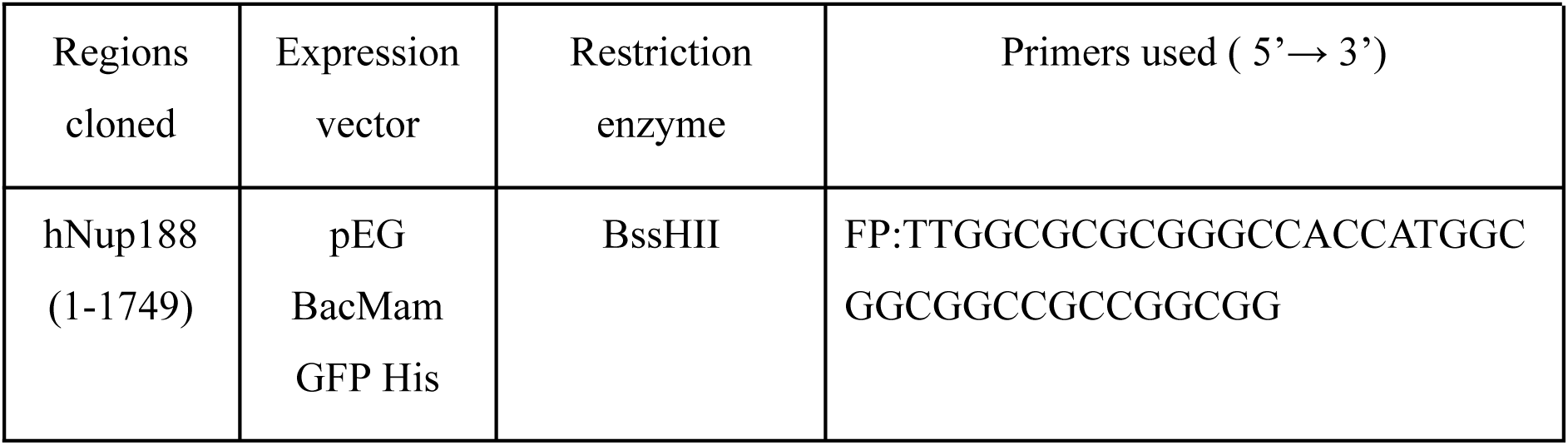
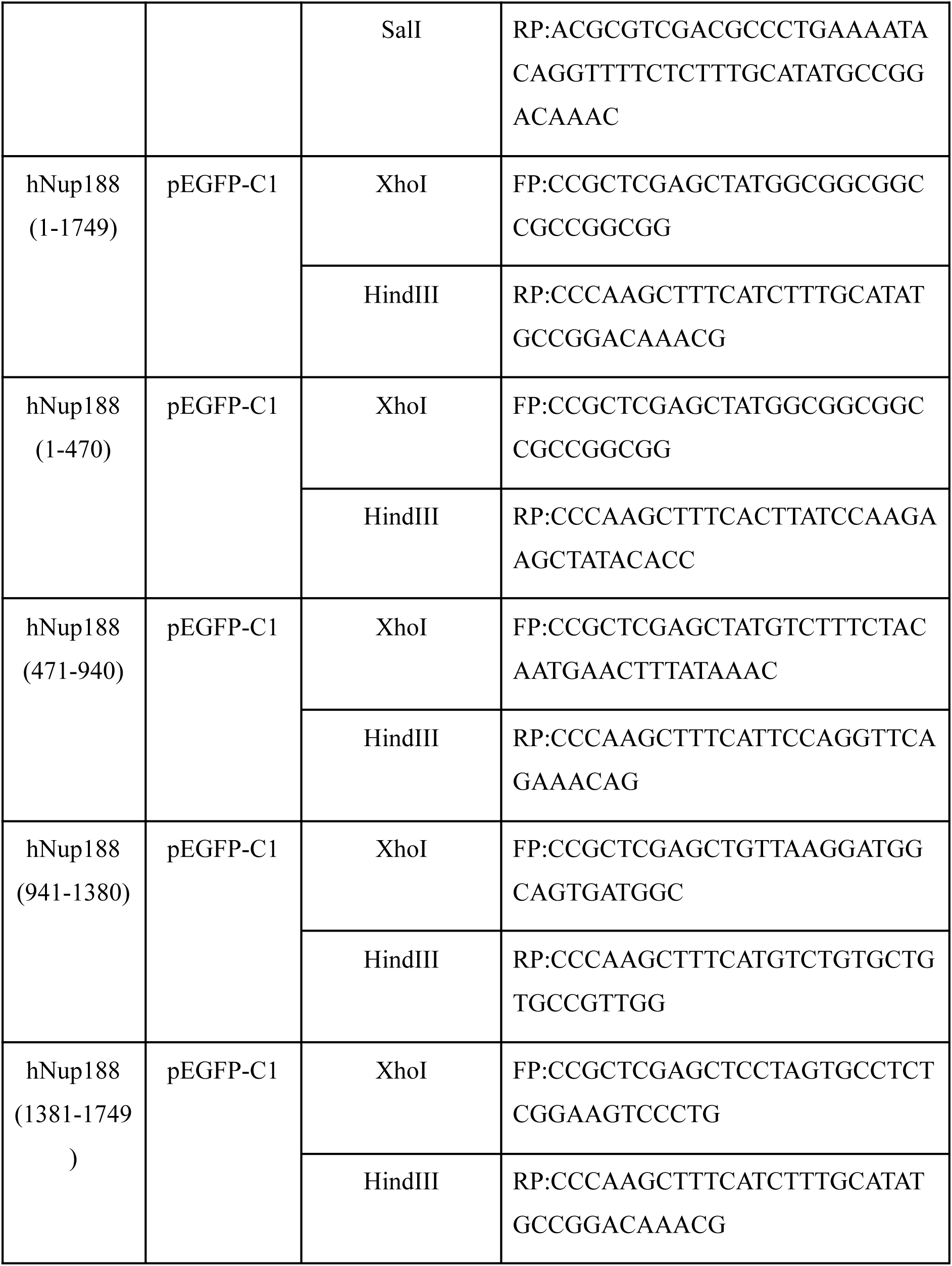

## Acknowledgments

We are grateful to Dr Deepa Subramanyam for sharing reagents. We thank the NCCS Bio-Imaging facility, and staff for their help with confocal-SIM microscopy. This work was supported by NCCS intramural funds and DBT grants (DBT/PR/26398 and BT/PR15450/COE/34/46/2016) awarded to R.C.. A.L.B. thanked DBT (DBT/2021-22/NCCS/1552) for the Ph.D. research fellowship. We thank members of the Lab of Structural Biology group for various suggestions and for critically reading the manuscript.

## Author Contributions

Conceptualization, Resources, Data curation, Formal analysis, Supervision, Funding acquisition, Validation, Investigation, Methodology, Writing-original draft, Project administration, Writing, review and editing - R.C.; Conceptualization, Data curation, Formal analysis; Validation; Investigation; Visualization; Methodology; Writing-original draft; Writing-review and editing - A.L.B.; Construct generation, Expression and purification of Nup188, Immunoprecipitation methodology, execution and analysis - B.B.; Construct generation, Optimization of purification of Nup188, Immunoprecipitation methodology, execution and analysis - S.B. MD Simulation methodology, analyzing, validation, Writing-original draft, review and editing, - S.D; MD Simulation execution and analysis - S.A.; Flickerprint execution and analysis, Reviewing - C.M.J.; Fibroblast culture, Immunofluorescence assay and analysis, reviewing - A.B.C.; Supervision - V.T.; Interactome of Nup188, reviewing and editing - K.R.; Fast paced Live recording of cells for Flickerprint - A.T.; Nup205 construct designing - P.S.; MD Simulation Supervision - S.C.M.; Conceptualization, Resources, Validation - J.J.; Flickerprint methodology and Supervision, review and editing - S.N.G.

## Declaration of generative AI and AI-assisted technologies in the writing process

During the preparation of this work, the author(s) used some portions of the language editing and initial drafting by Claude AI and Google Gemini. All content was subsequently reviewed and verified by the authors. After using this tool/service, the author(s) reviewed and edited the content as needed and take(s) full responsibility for the content of the published article.

## Contact for reagent and resource sharing

Further information and requests for resources and reagents should be directed to and will be fulfilled by the Lead Contact Radha Chauhan.

## Supplemental Information

Supplemental information includes 17 figures, 6 Movie files and 1 excel file.

**Figure S1.**
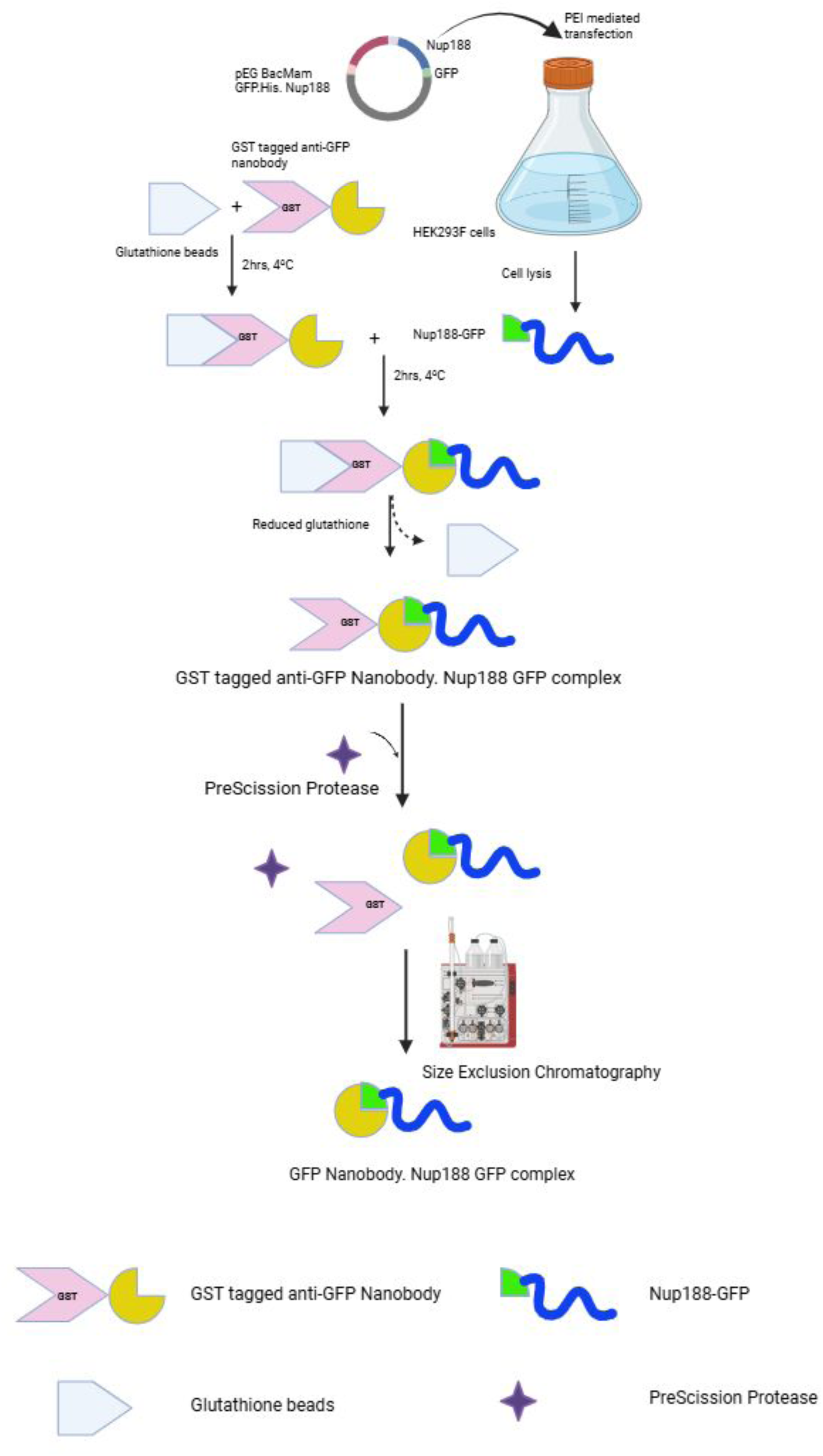
Purification of human Nup188 from HEK293F cells by GFP-nanobody affinity capture. HEK293F cells were transfected with pEG BacMam GFP.His.Nup188 using PEI. GST-tagged anti-GFP nanobody was immobilized on glutathione beads and incubated with cell lysate to capture Nup188-GFP. The complex was eluted with reduced glutathione, yielding GST tagged GFP nanobody–Nup188-GFP. Subsequent removal of the GST tag by PreScission protease followed by SEC results in the pure form of the Nup188-GFP protein.

**Figure S2.**
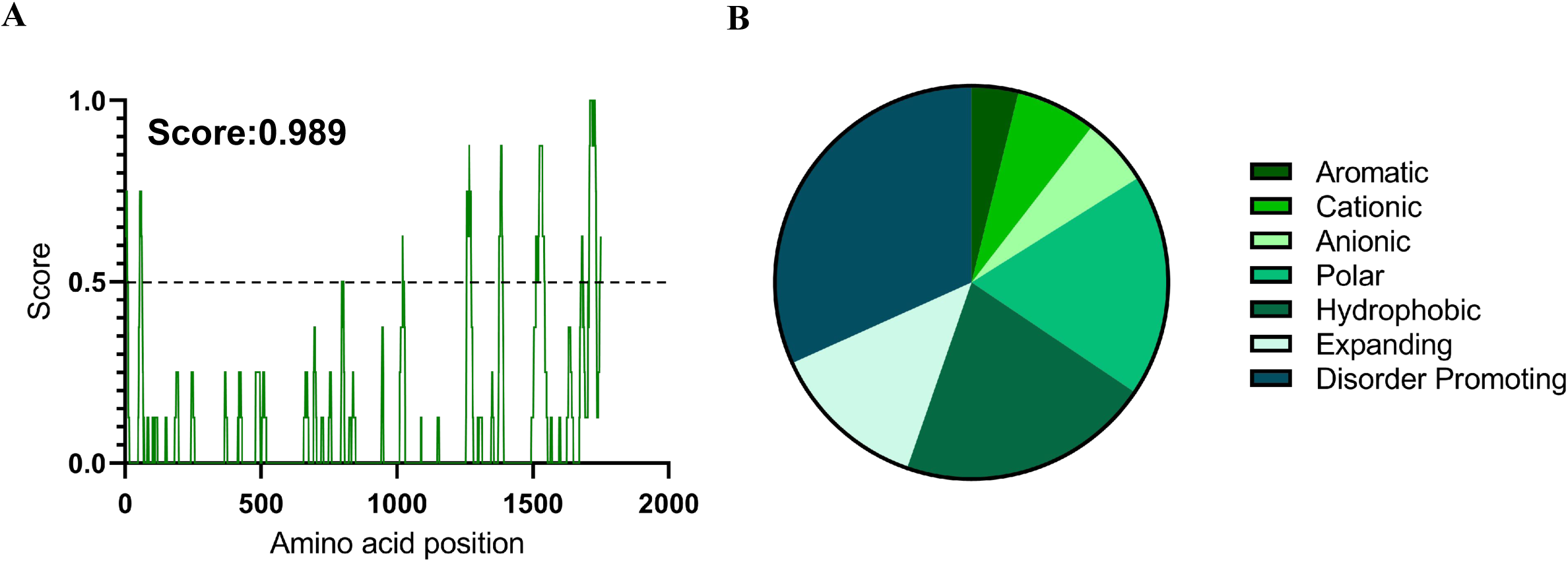
MolPhase analysis of human Nup188. (A) Per-residue MolPhase score along the Nup188 sequence. The dashed line marks the threshold (0.5); the overall score for Nup188 is 0.989. (B) Amino acid composition of Nup188 by residue class (aromatic, cationic, anionic, polar, hydrophobic, expanding and disorder-promoting), showing each class’s contribution to the predicted phase separation propensity.

**Figure S3.**
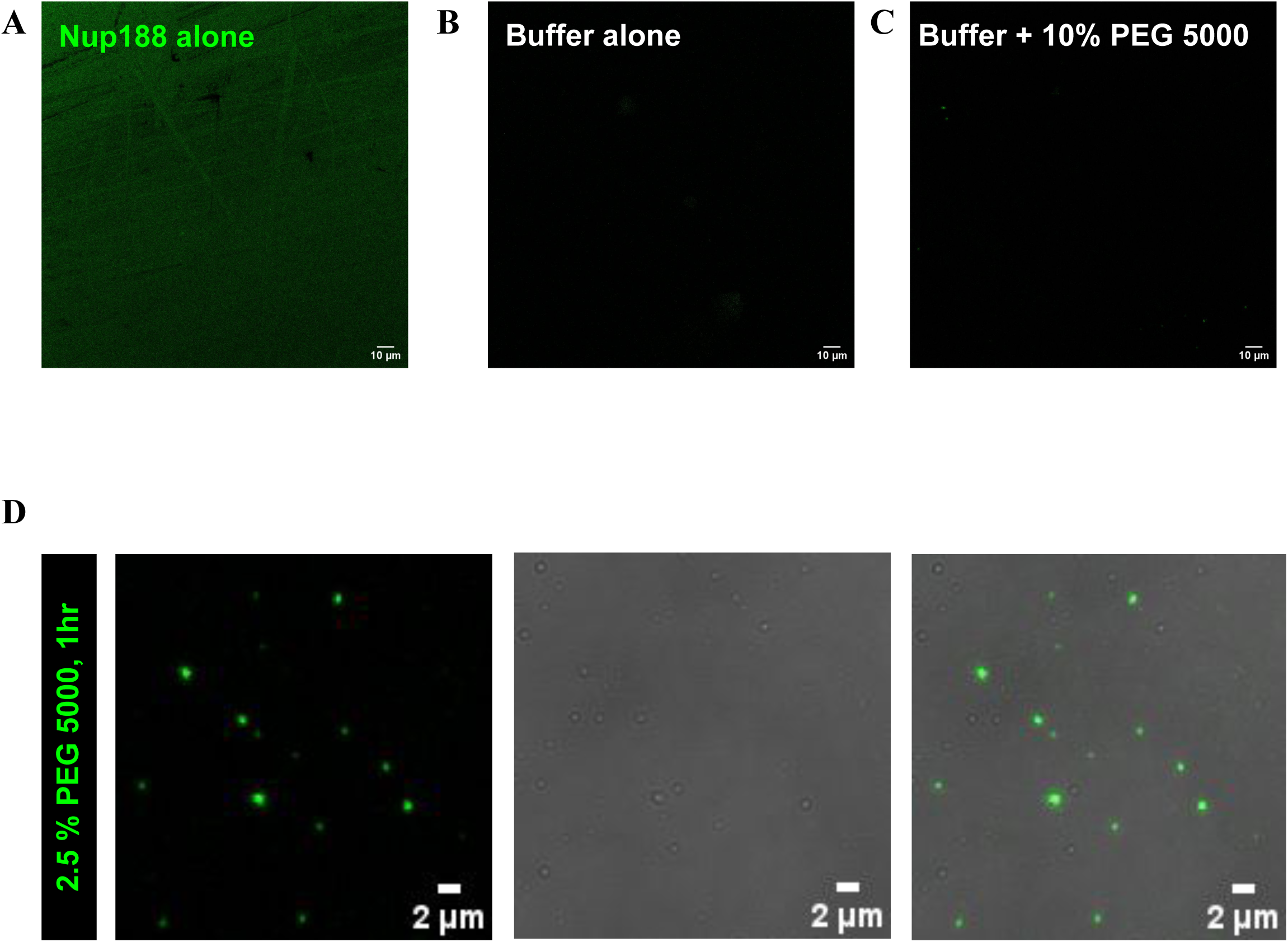
Controls and low-PEG condensate formation for recombinant Nup188. (A–C) Fluorescence images of (A) purified Nup188 after high-speed centrifugation without PEG, (B) buffer alone and (C) buffer with 10 % PEG 5000. None show condensates. (D) Fluorescence, DIC and merged images of Nup188 with 2.5 % PEG 5000 after 1 h, showing condensate formation. Scale bars, 2 µm. Representative of three independent experiments.

**Figure S4.**
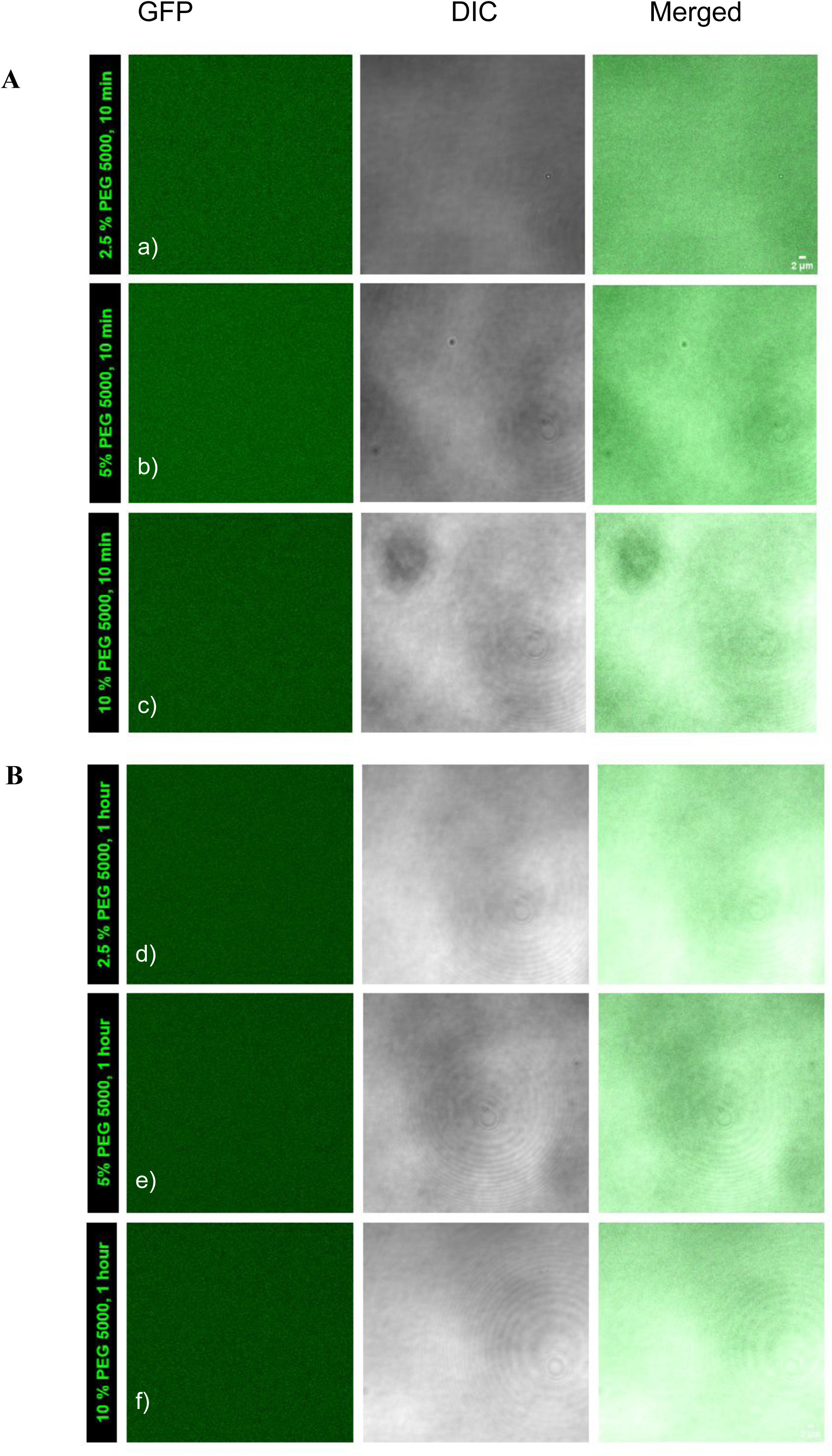
Recombinant GFP does not form condensates in DDM-containing buffers. GFP, DIC and merged images of recombinant GFP mixed with PEG 5000 at 2.5 % (a, d), 5 % (b, e) or 10 % (c, f), imaged after (A) 10 min or (B) 1 h. No condensates form under any condition. Scale bar, 10 µm. Representative of three independent experiments.

**Figure S5.**
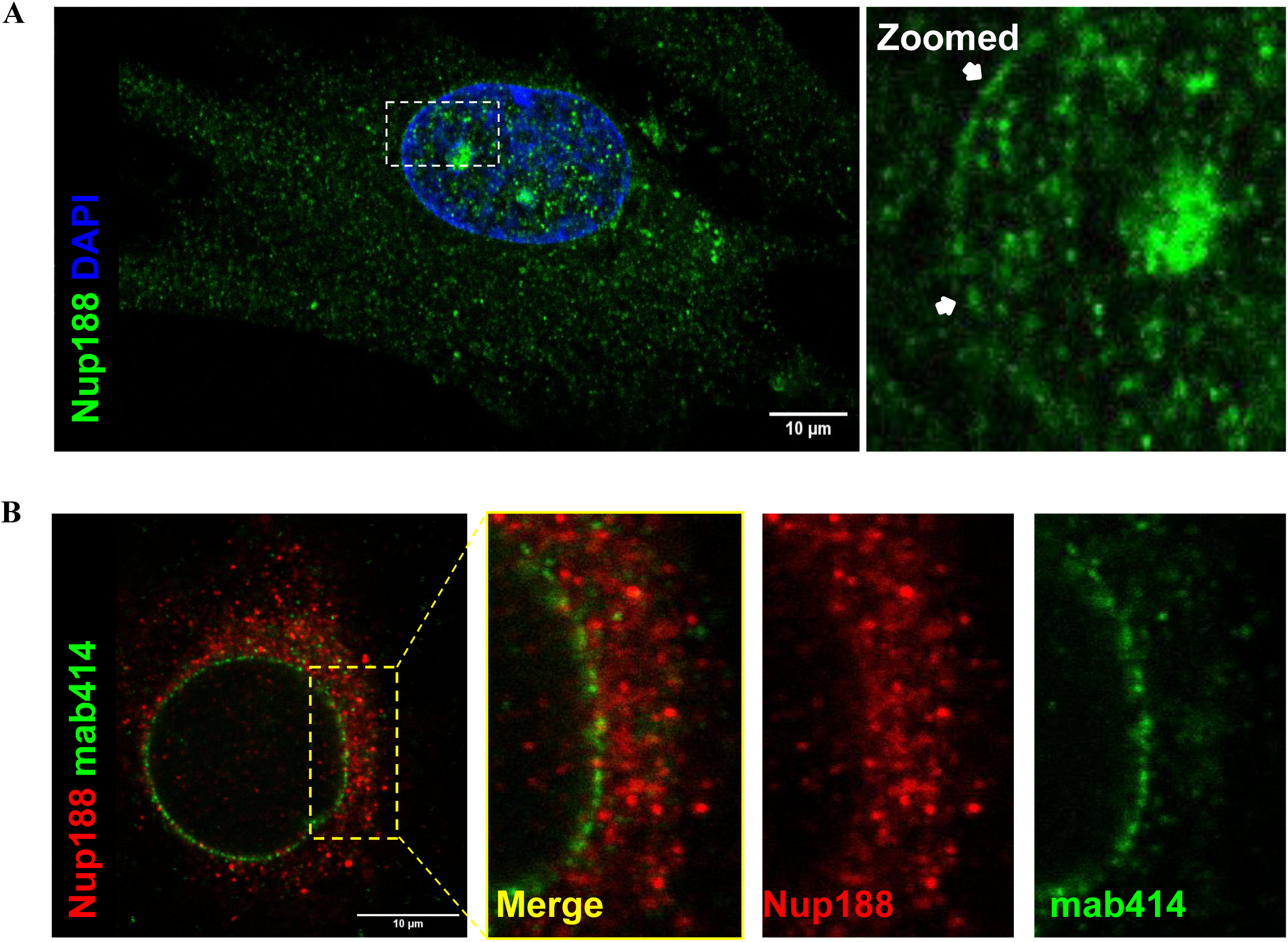
Endogenous Nup188 localization. (A) Immunofluorescence of endogenous Nup188 (green) and DAPI (blue) in WI-38 cells; this is the image shown in Fig. 3A, reproduced here to highlight nuclear envelope staining (arrows in the enlarged inset). (B) Co-staining of endogenous Nup188 (red) and mab414 (nuclear pore complexes, green) in U2OS cells; merged and single channels of the boxed region are shown on the right. Scale bars, 10 µm. Three independent experiments, 10 cells each.

**Figure S6.**
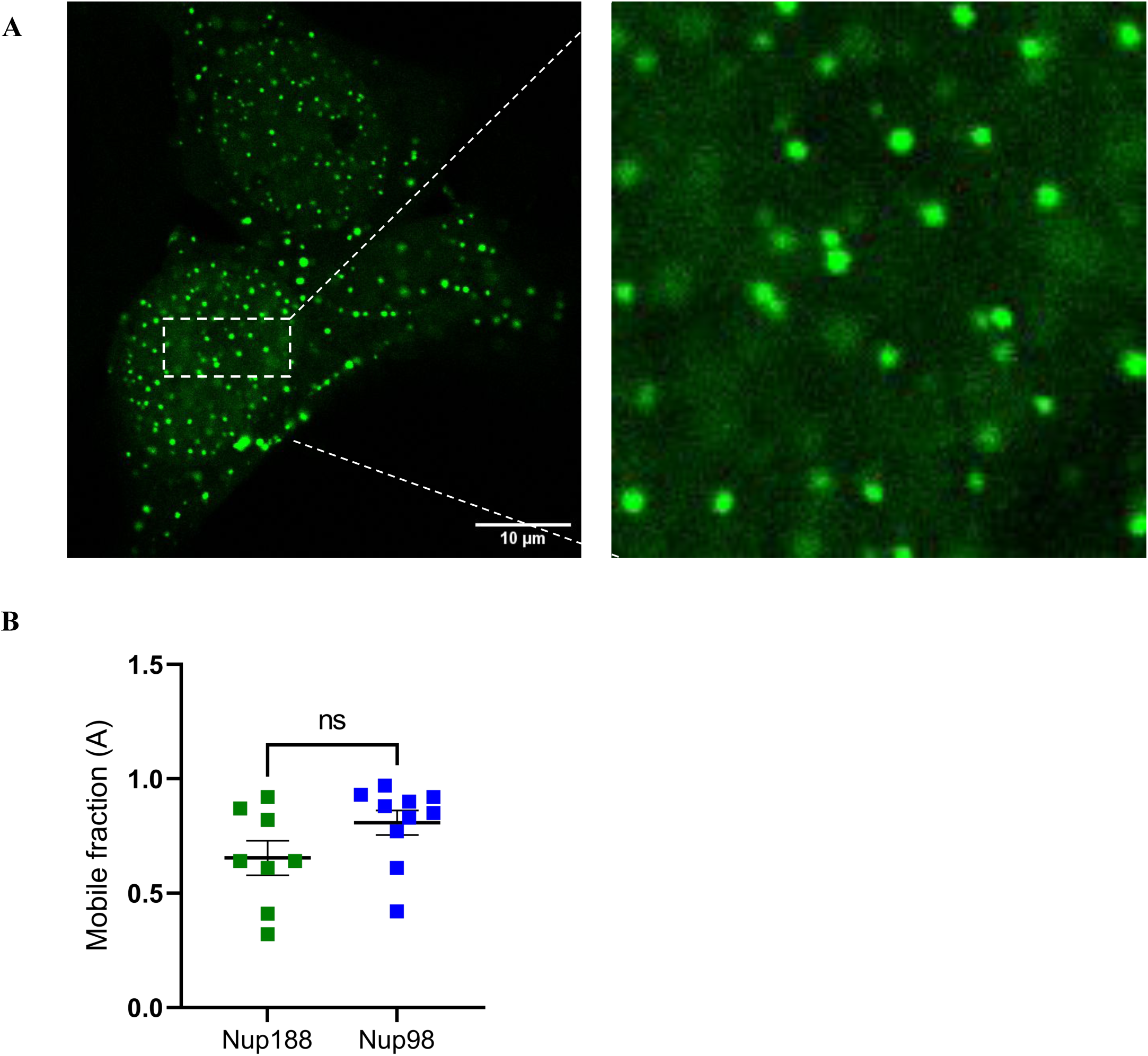
Nup188 forms condensates in HEK293FT cells. (A) Confocal image of HEK293FT cells expressing Nup188-GFP, which forms nuclear and cytoplasmic condensates (boxed region enlarged on the right). Scale bar, 10 µm. N = 3 experiments, n = 10 cells each. (B) FRAP mobile fraction (A) of Nup188 and Nup98 condensates from U2OS cells, with Nup98 as a reference phase-separating FG-nucleoporin. Each square is one condensate; horizontal bar, mean ± SEM. ns, not significant (p = 0.0962; two-tailed Mann–Whitney U test).

**Figure S7.**
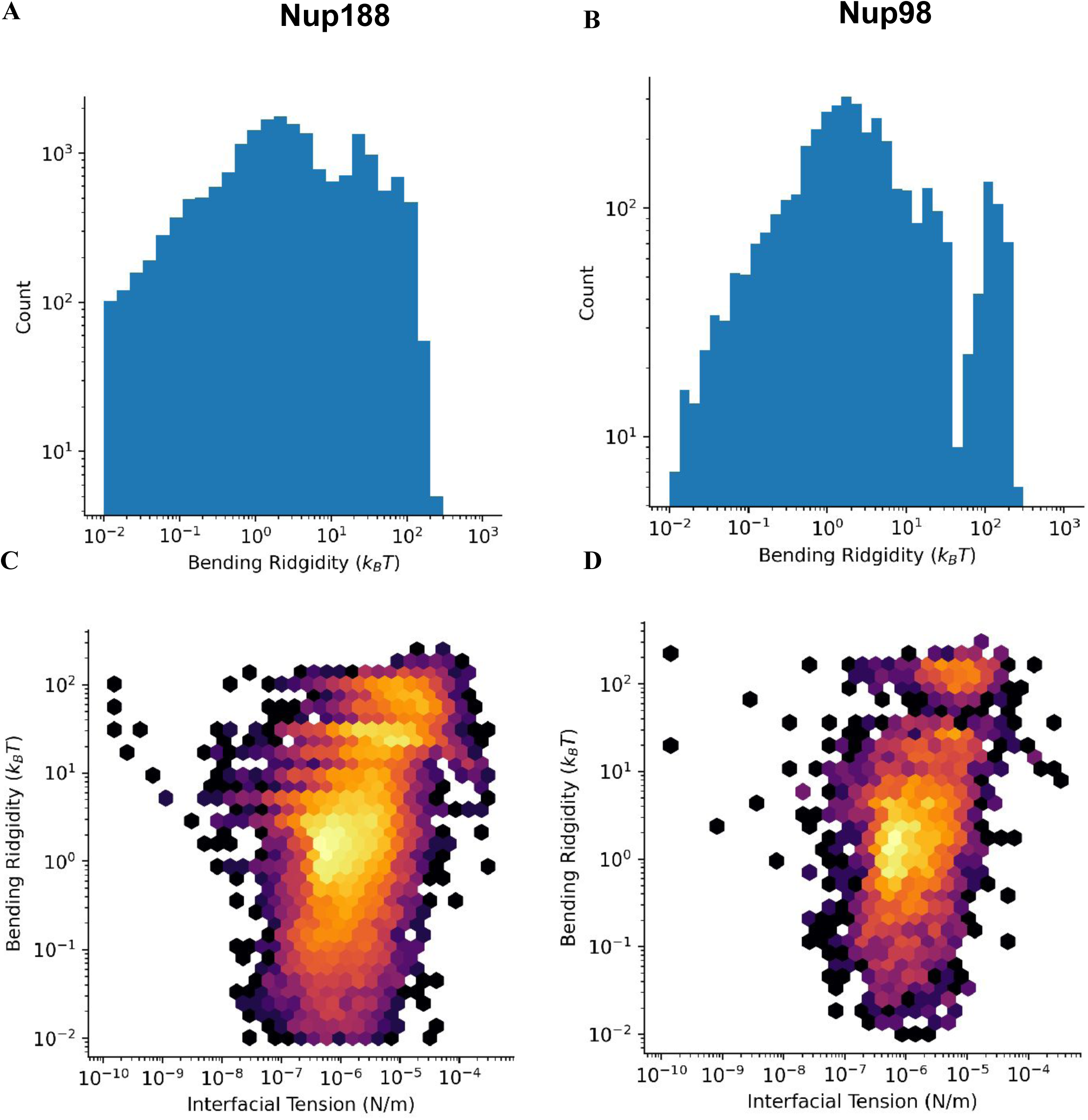
Relationship between bending rigidity and interfacial tension of Nup188 and Nup98 condensates. FlickerPrint analysis. (A, B) Histograms of bending rigidity (κ) for Nup188 (A) and Nup98 (B) condensates. (C, D) Density scatter plots of κ against interfacial tension (σ) for Nup188 (C) and Nup98 (D). Colour indicates local point density (yellow, high; purple, low). The main population of both condensate types occupies a similar κ–σ range.

**Figure S8.**
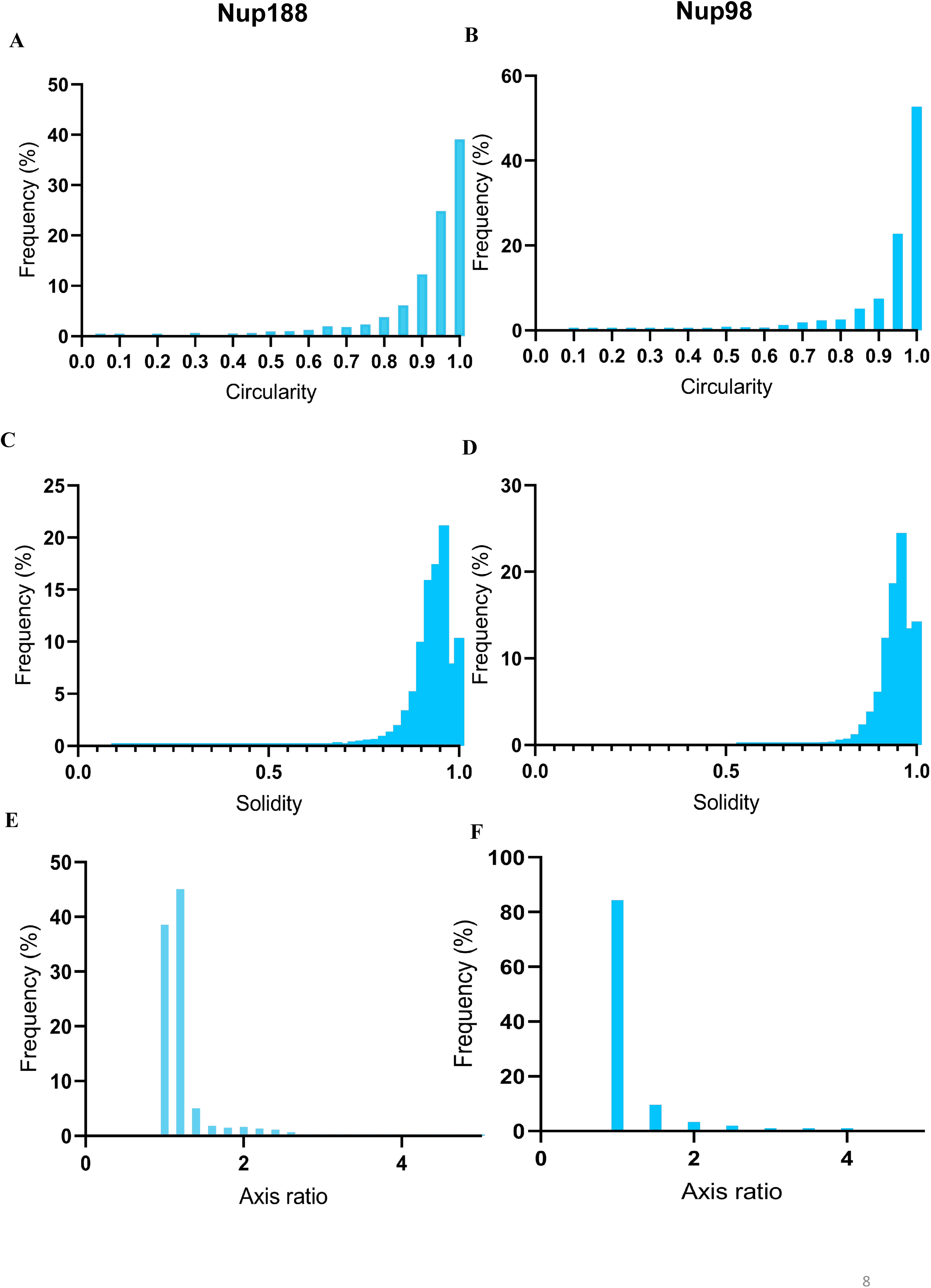
Shape descriptors of Nup188 and Nup98 condensates from FlickerPrint. Frequency distributions of (A, B) circularity, (C, D) solidity and (E, F) axis ratio for Nup188 (A, C, E) and Nup98 (B, D, F) condensates. Values near 1 indicate near-spherical condensates.

**Figure S9.**
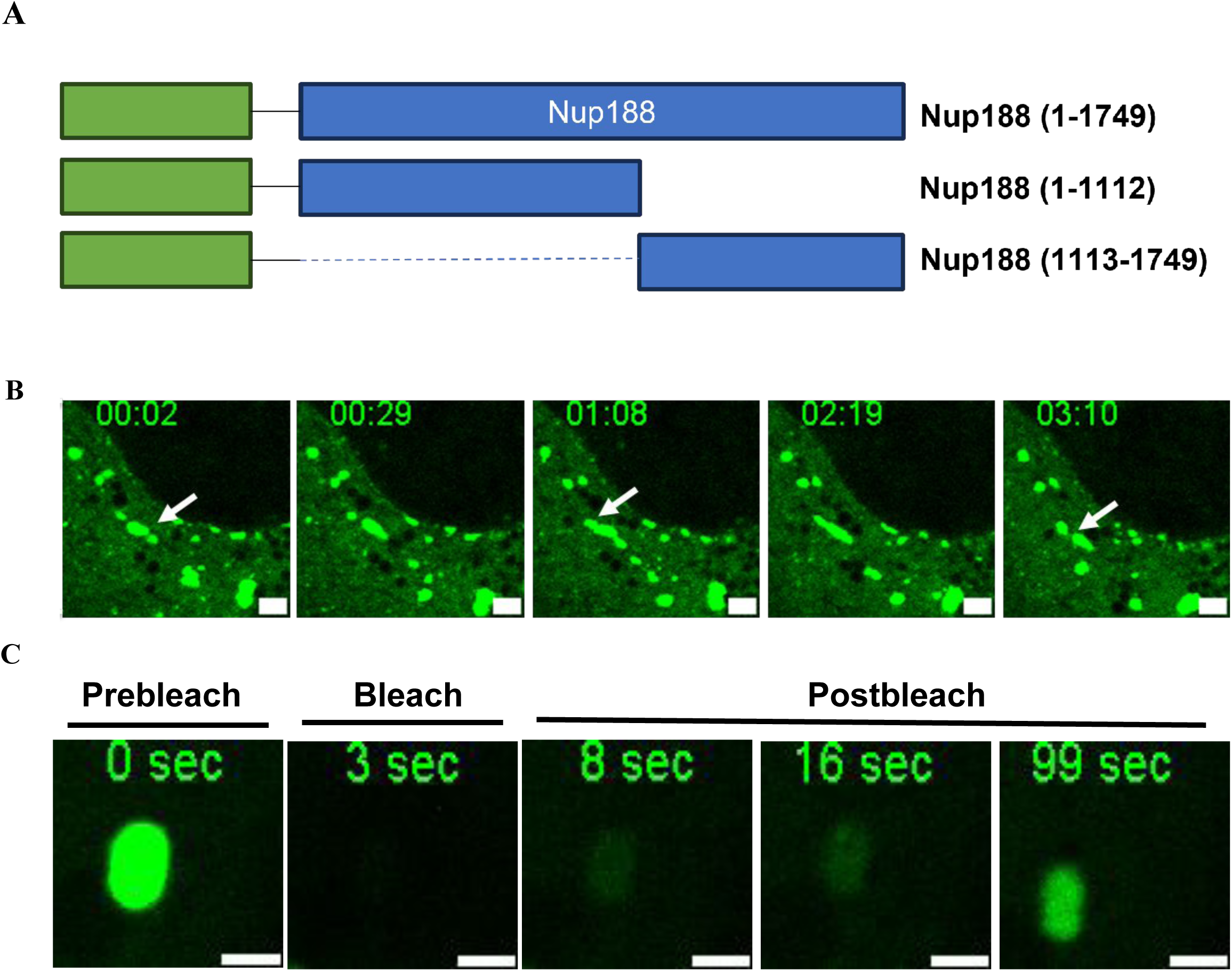
The N- and C-terminal fragments of Nup188 differ in condensate formation and dynamics. (A) Nup188 construct used: Full length (1-1749), N-terminal fragment (1-1112) and C-terminal fragment (1113-1749) (B) Time-lapse of condensates in a live cell (time in mm:ss); arrows follow fusion-fission events. (C) FRAP of a single condensate before bleaching, at bleaching and post bleaching. Scale bars 2 µm

**Figure S10.**
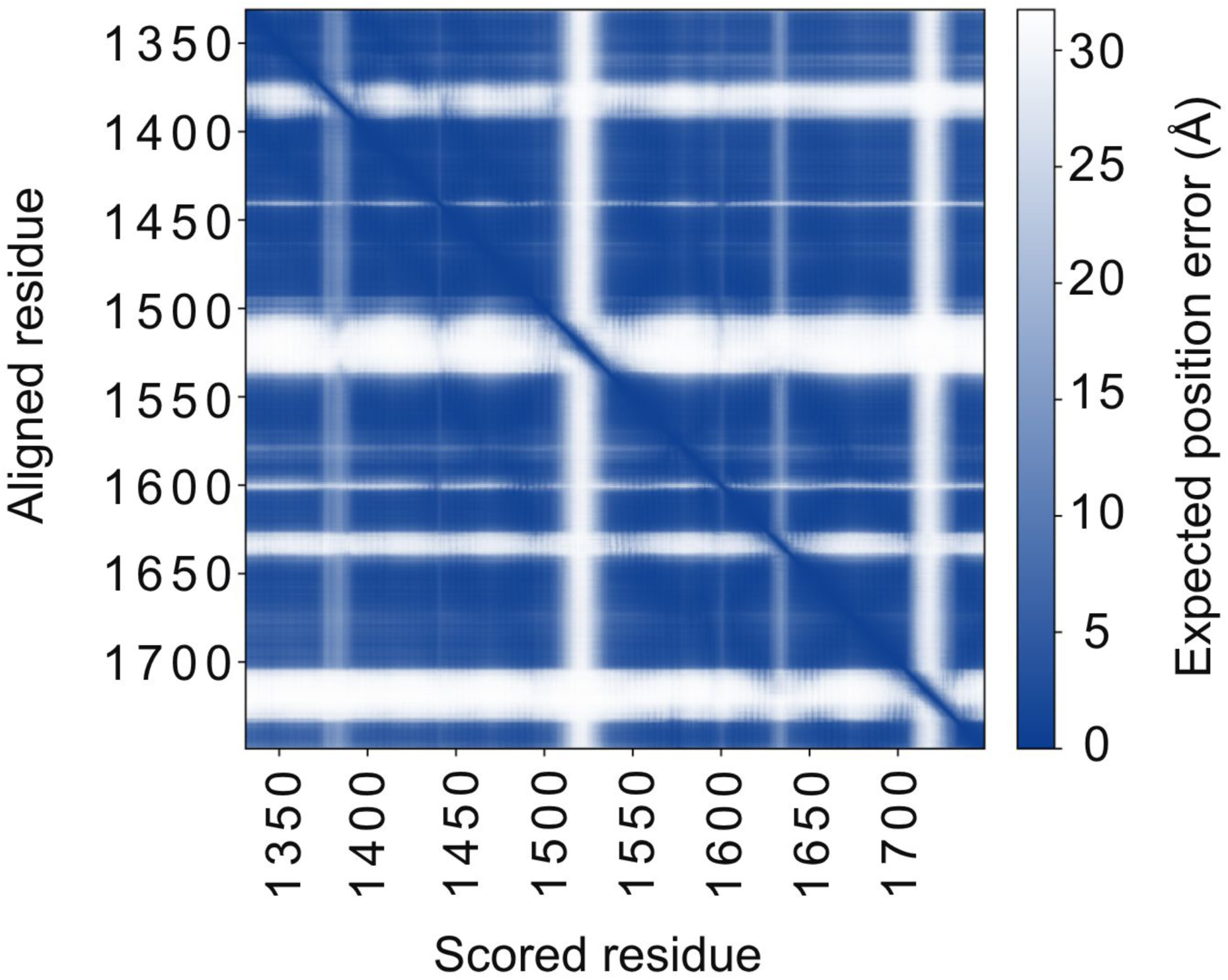
Predicted aligned error (PAE) of the AlphaFold 3 model of Nup188(1331–1749) region. Each cell shows the expected position error (Å) of the scored residue (x-axis) when the model is aligned on the aligned residue (y-axis). Dark blue indicates low error (high confidence) and white indicates high error (low confidence). The dark diagonal shows that local backbone geometry is confidently predicted, and the dark off-diagonal blocks indicate confidently predicted substructures. Three prominent white bands at ∼1370–1390, ∼1500–1540 and ∼1700–1745, with thinner bands at ∼1440, ∼1600 and ∼1635, indicate uncertain relative orientation of the segments on either side, consistent with flexible linkers or hinges. The three major bands largely coincide with IDR1-3, identified by MD simulations (Fig 8D)

**Figure S11.**
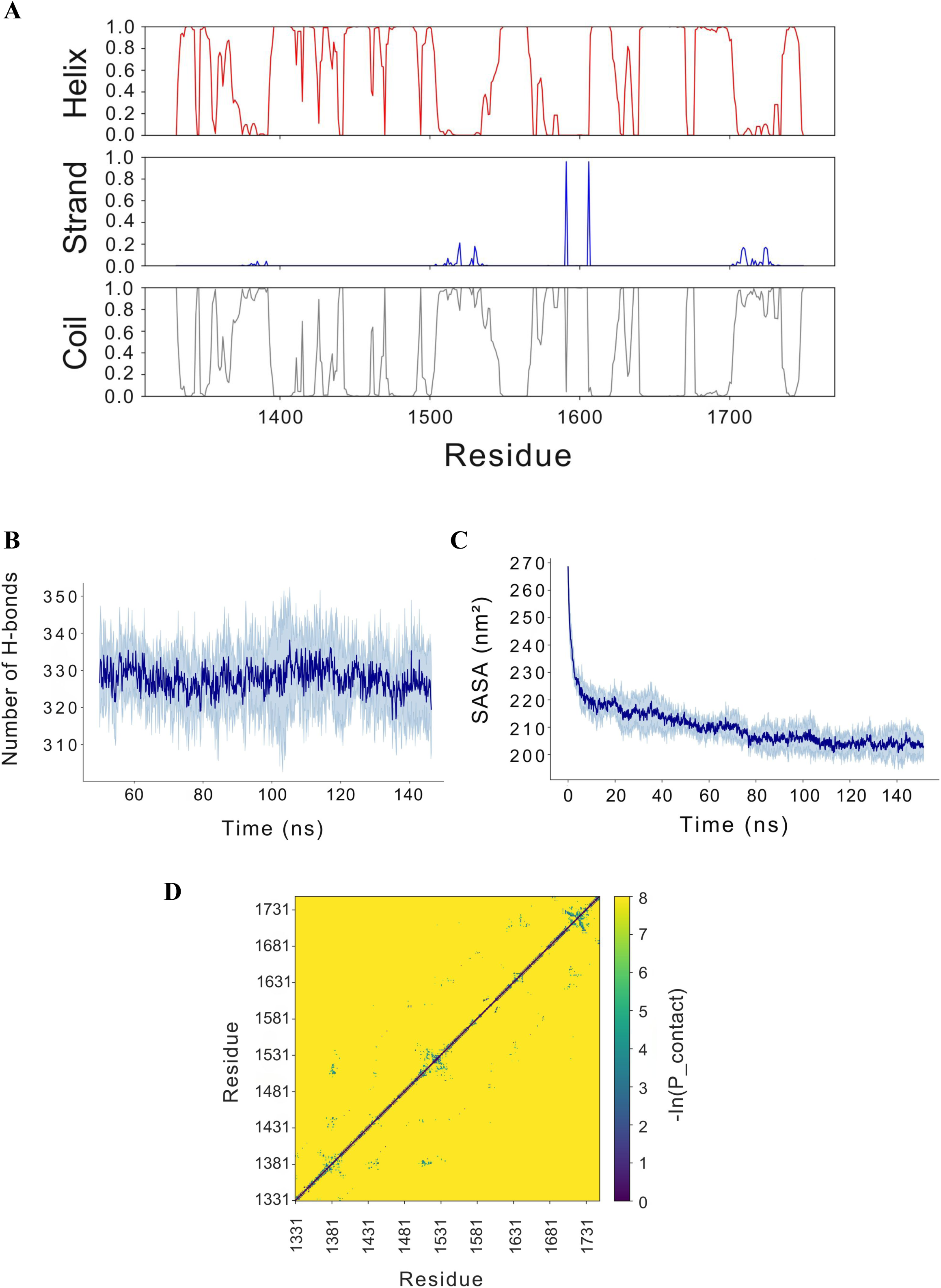
All-atom simulations of the Nup188 C-terminus (CHARMM36m, six replicas). (A)Per-residue secondary structure probability is shown from DSSP averaged over replicas: helix (top, red), β-strand (middle, blue), coil (bottom, grey). (B) Number of intramolecular hydrogen bonds over the equilibrated portion of the trajectories, plateauing at ∼328. (C) Solvent-accessible surface area (SASA). In B and C the dark line is the replica mean and shading ±1 SD. (D) Intra-chain residue–residue contact map plotted as −ln(P_contact); dark blue/purple: frequent contacts; and yellow color represent no contacts. Contacts are dense along the diagonal and sparse at long range.

**Figure S12.**
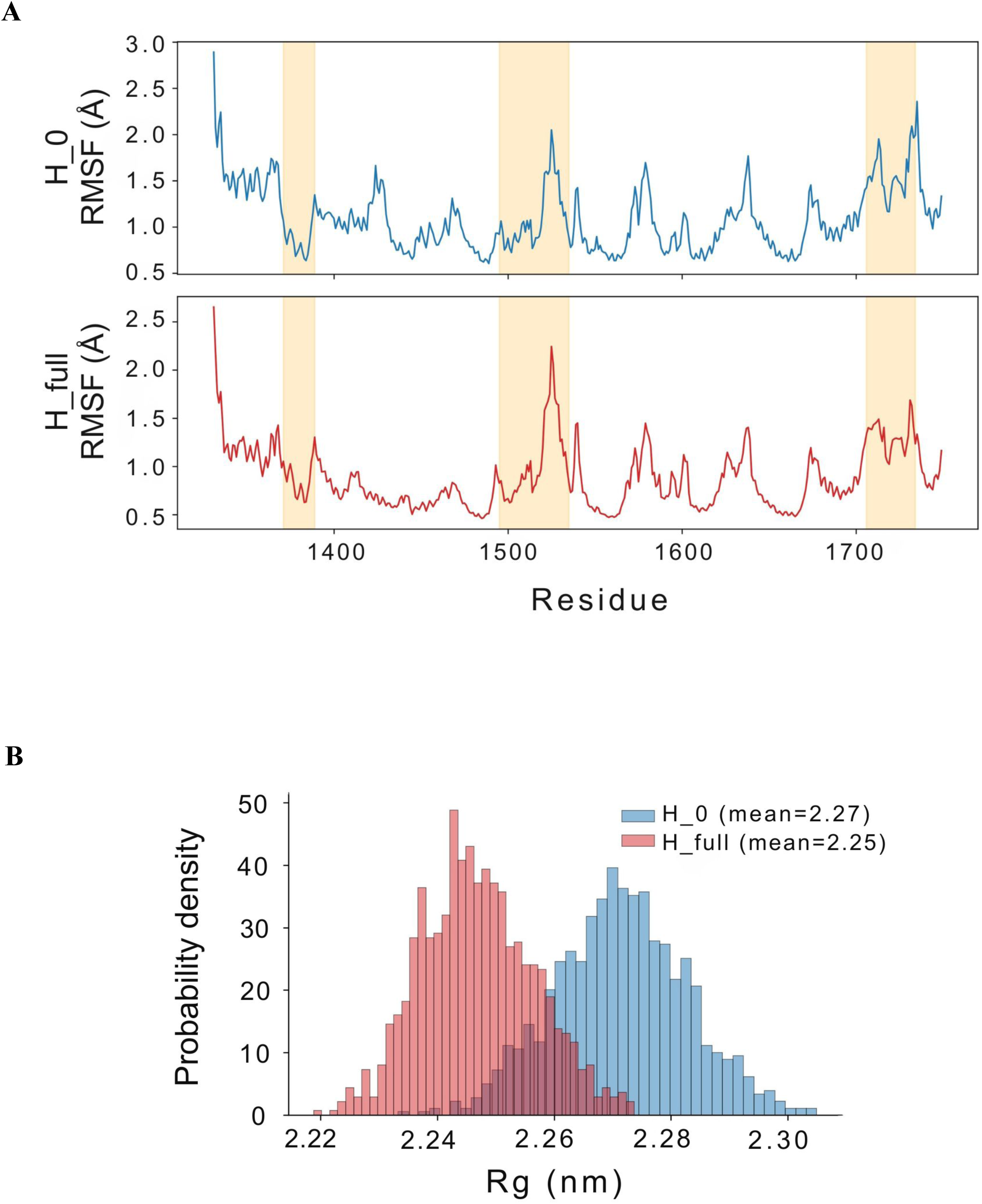
Coarse-grained (Martini 3) single-chain simulations of the Nup188 C-terminus. (A) Per-residue RMSF for H_0 (all-coil, top, blue) and H_full (DSSP-derived helices retained, bottom, red). Shaded bands are the predicted IDRs. (B) Distributions of single-chain radius of gyration (Rg) at 310 K for H_0 (blue; mean 2.27 nm) and H_full (red; mean 2.25 nm).

**Figure S13.**
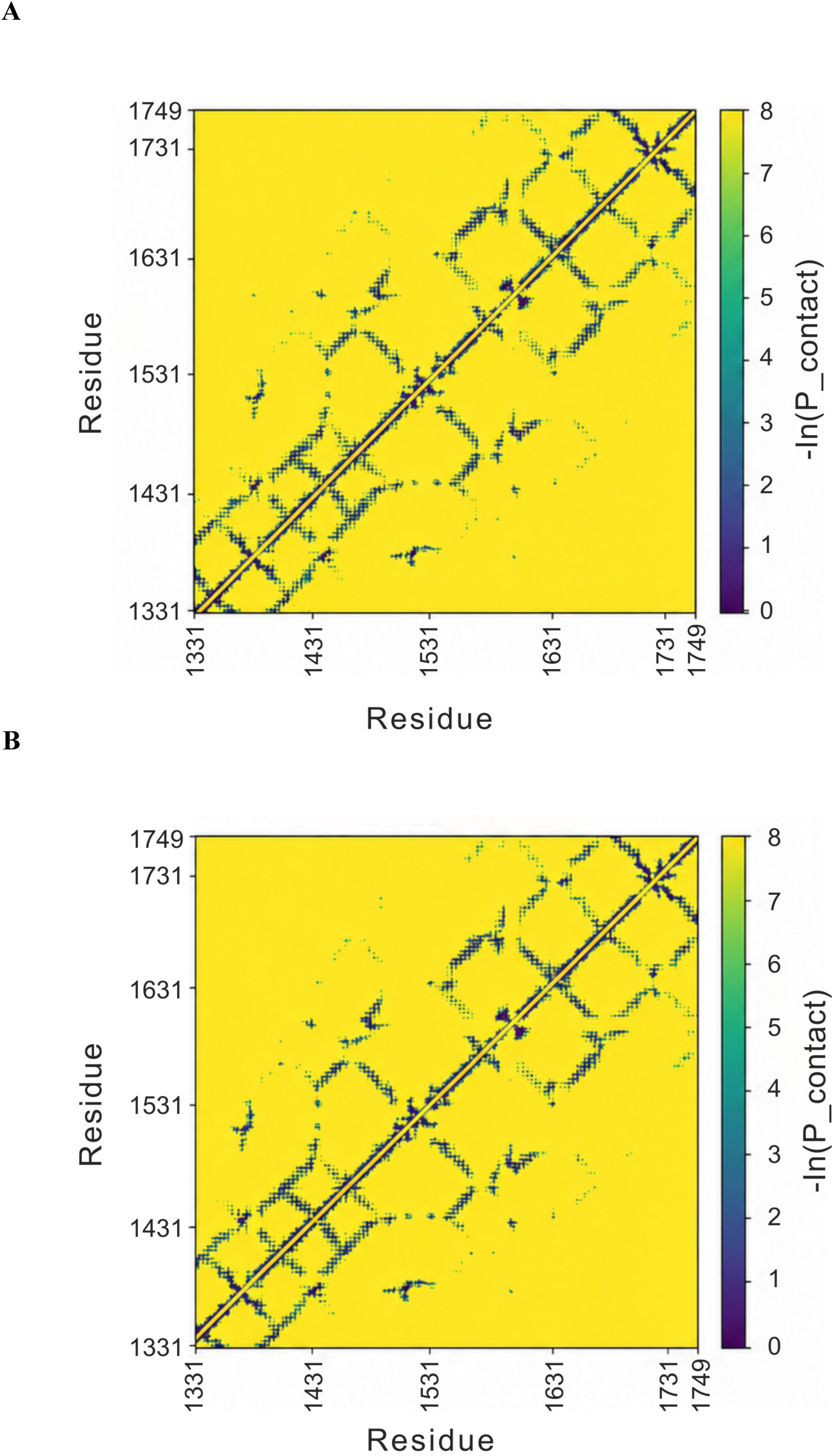
Intra-chain contact maps of the Nup188 C-terminus from coarse-grained simulations. Residue–residue contact maps (−ln(P_contact)) for H_0 (top) and H_full (bottom) are shown over 1 µs of Martini 3 simulation. Dark blue indicates frequent contacts, yellow indicates none.

**Figure S14.**
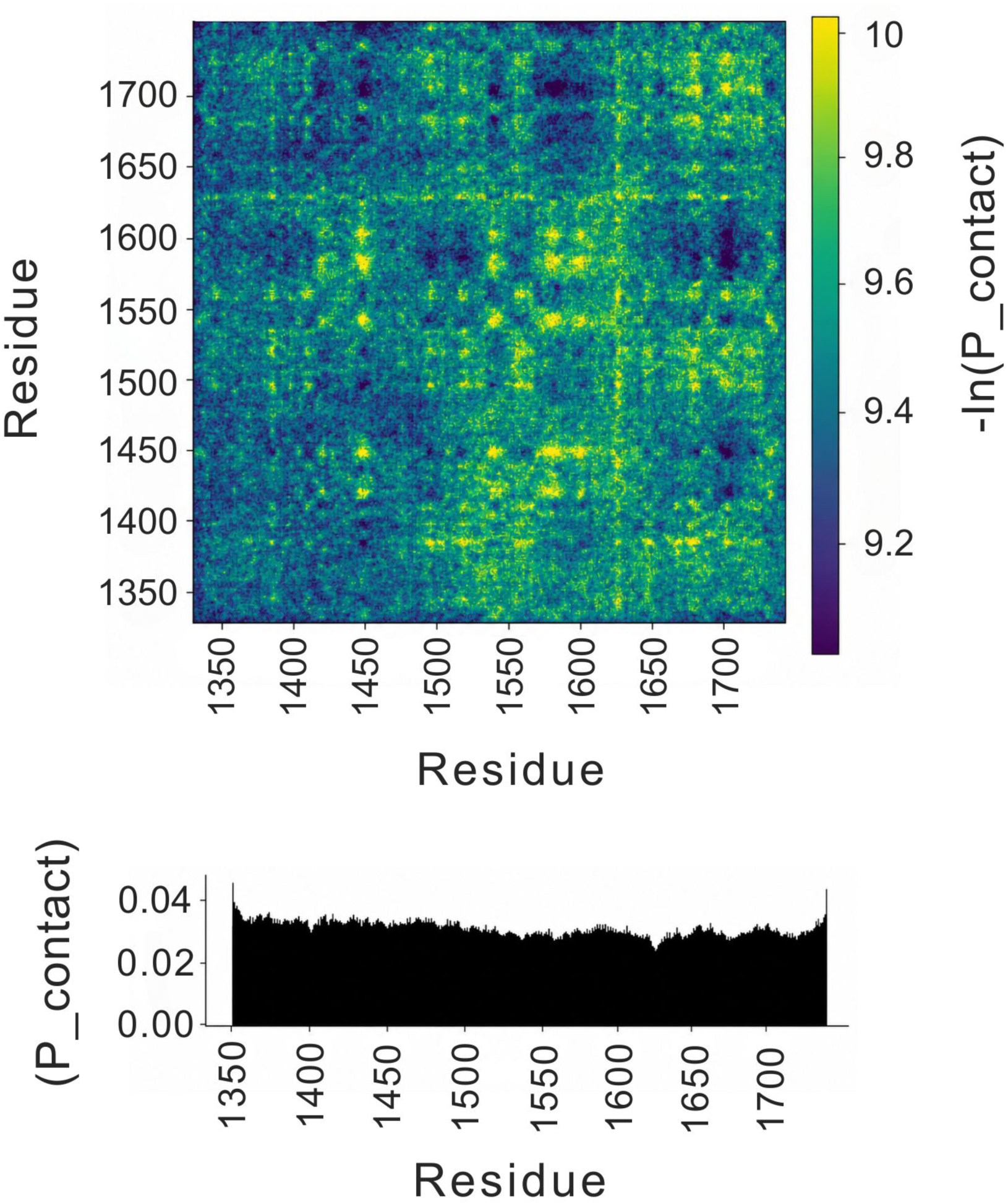
Intermolecular contact map from the HPS-Urry slab simulation. Inter-chain residue–residue contact probability is plotted as −ln(P_contact), with the colour scale stretched to the populated range (∼9–10). Contacts are diffuse along the sequence rather than concentrated at discrete hot spots, unlike the site-specific pattern in the Martini simulations (Figure 9B). The lower panel shows per-residue contact probability, which is approximately uniform.

**Figure S15.**
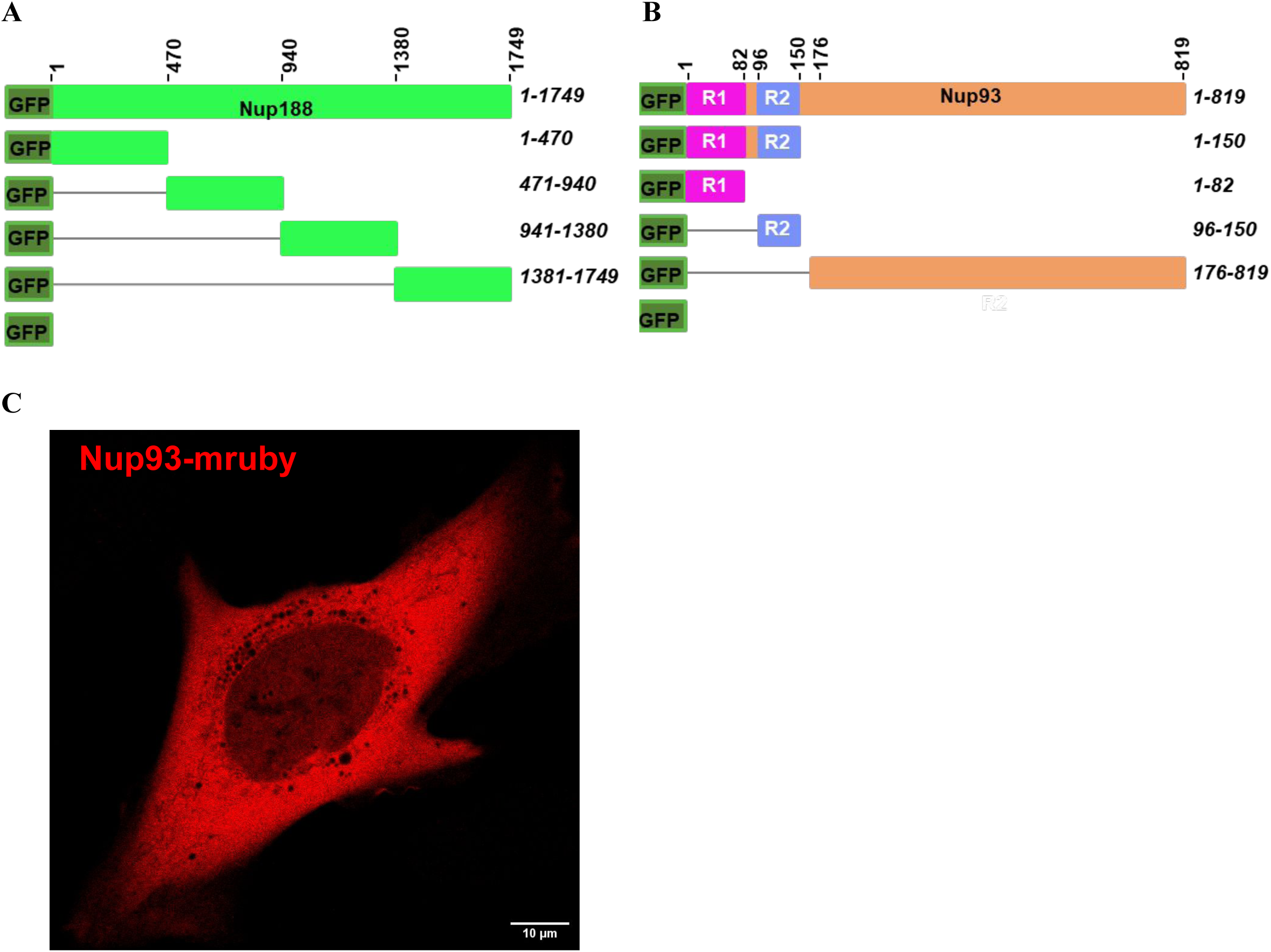
Constructs used in this study for mapping the Nup93–Nup188 interaction. (A) GFP-Nup188 constructs: 1–1749, 1–470, 471–940, 941–1380, 1381–1749 and GFP alone. (B) GFP-Nup93 constructs used in pull-downs: 1–819, 1–150, 1–82 (R1), 96–150 (R2), 176–819 and GFP alone. (C) U2OS cell expressing Nup93-mRuby, which is diffusely distributed. Scale bar, 10 µm.

**Figure S16.**
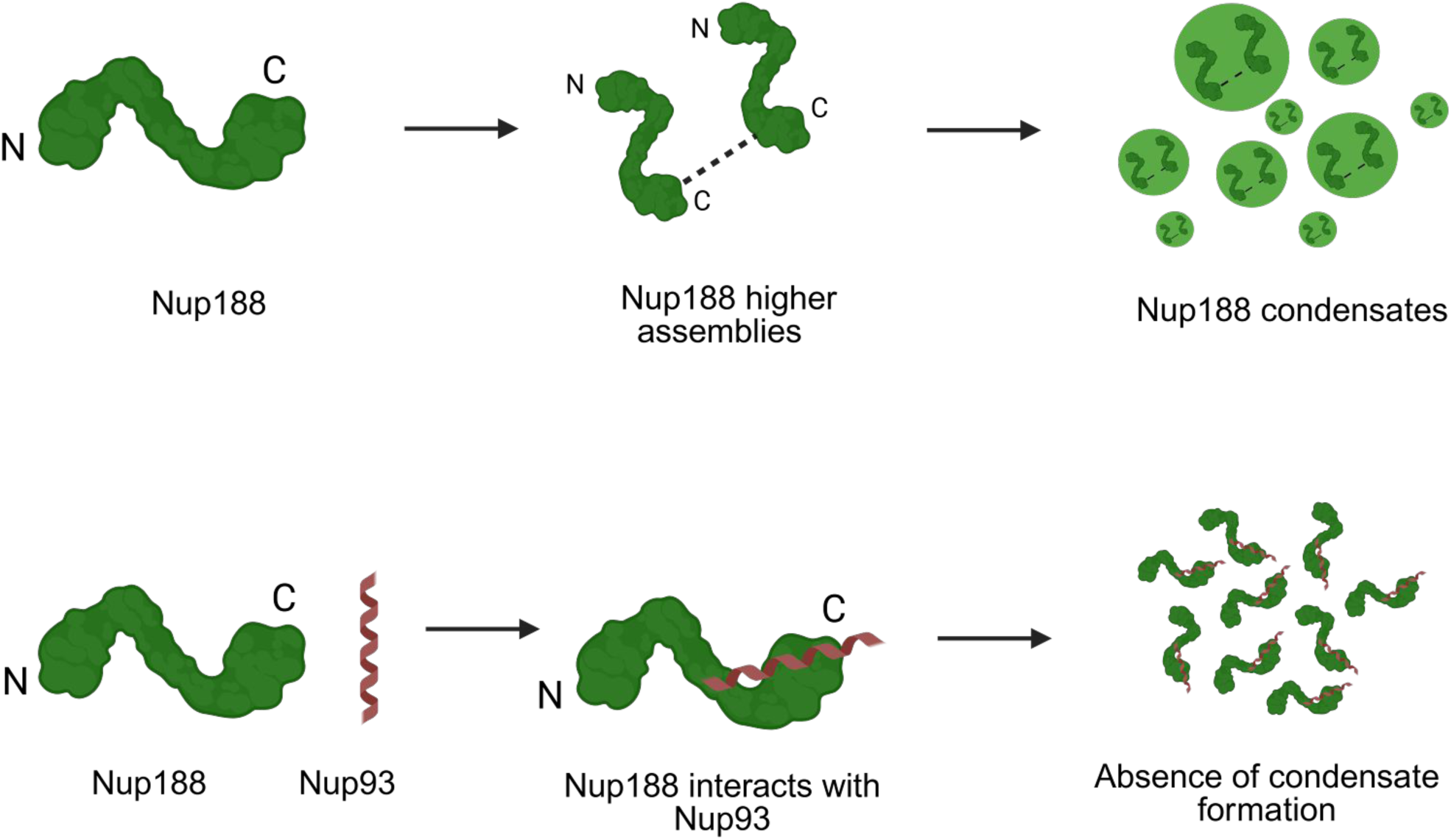
Model for disruption of Nup188 condensates by Nup93. Top: Nup188 forms higher-order assemblies through C-terminal interactions and then condensates. Bottom: Binding of Nup93 to the Nup188 C-terminal region prevents this assembly, so condensates do not form. N and C mark the termini.

**Figure S17.**
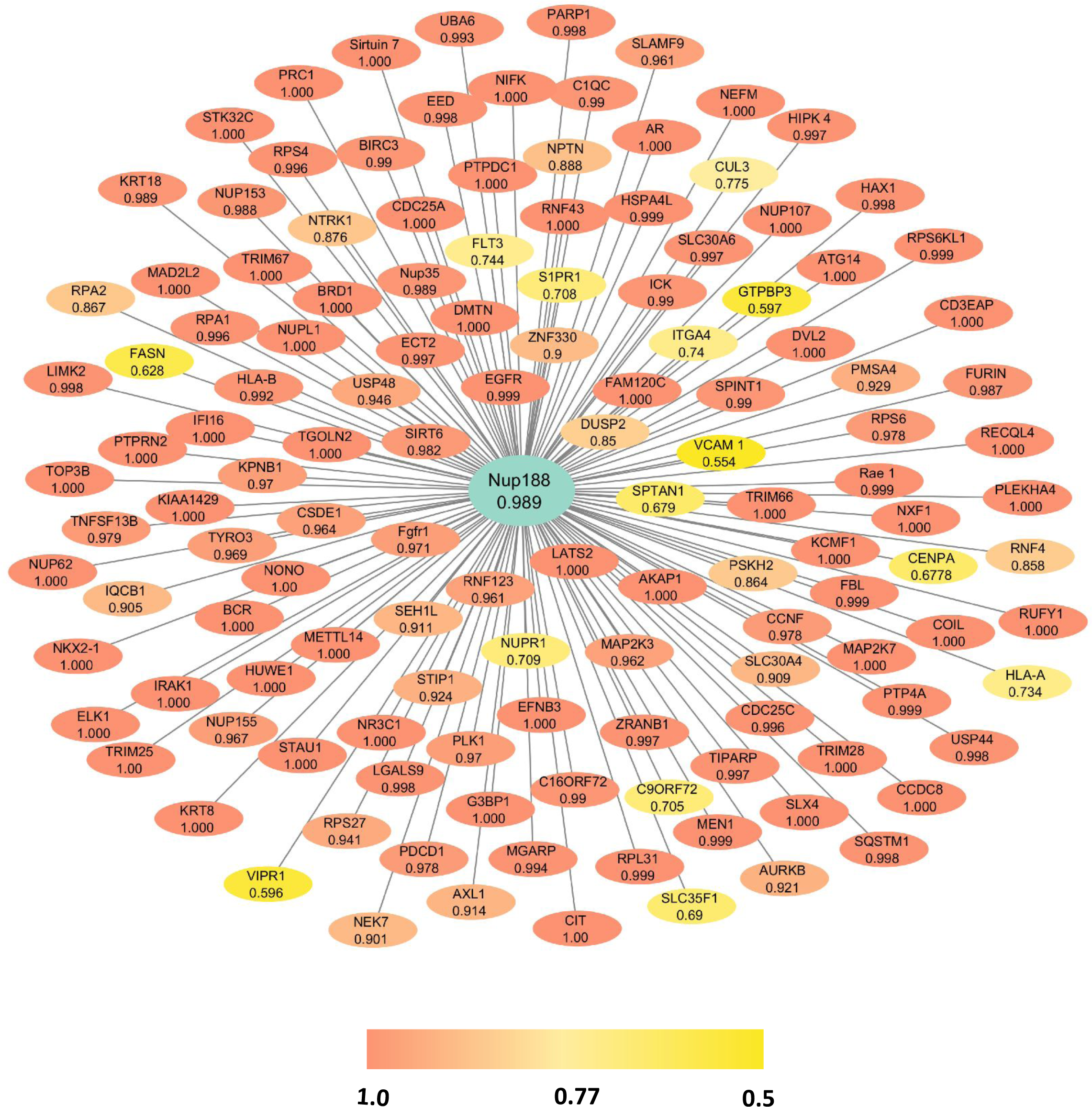
Protein–protein interaction network of Nup188 with MolPhase scores. BioGRID interactors of human Nup188 (centre node, teal) from high- and low-throughput studies, visualized in Cytoscape. Only interactors with a MolPhase LLPS score > 0.5 are shown. The number in each node is its MolPhase score. Node colour follows the score: yellow (0.5–0.65), light orange (0.65–0.8) and dark orange (0.8–1.0).

## Supplementary video legends

**Supplementary Video 1: Live imaging of Nup188 expressing U2OS cells**

In video, 1 sec = 20 sec

**Supplementary Video 2: Fusion of Nup188 condensates in U2OS cells**

1 sec = 10 sec

**Supplementary Video 3: Dynamics of Nup188 condensates in U2OS cells**

**Supplementary Video 4: 2D projection of immunostaining of Nup188GFP cells with Nup358 antibodies**

**Supplementary Video 5: Nup188 24-mer assembly in MD**

**Supplementary Video 6: Live imaging of Co-expression of Nup188 and Nup93**

**Supplementary files**

**Supplementary file 1: Excel sheet representing Protein-protein interaction network of human Nup188 obtained from BioGRID with MolPhase LLPS propensity score.**

